# Integrative Clinical and Molecular Characterization of Translocation Renal Cell Carcinoma

**DOI:** 10.1101/2021.04.14.439908

**Authors:** Ziad Bakouny, Ananthan Sadagopan, Praful Ravi, Nebiyou Y. Metaferia, Jiao Li, Shatha AbuHammad, Stephen Tang, Thomas Denize, Emma R. Garner, Xin Gao, David A. Braun, Laure Hirsch, John A. Steinharter, Gabrielle Bouchard, Emily Walton, Destiny West, Chris Labaki, Shaan Dudani, Chun-Loo Gan, Vidyalakshmi Sethunath, Filipe LF. Carvalho, Alma Imamovic, Cora Ricker, Natalie I. Vokes, Jackson Nyman, Jihye Park, Michelle S. Hirsch, Rizwan Haq, Gwo-Shu Mary Lee, Bradley A. McGregor, Steven L. Chang, Adam S. Feldman, Catherine J. Wu, David F. McDermott, Daniel Y.C. Heng, Sabina Signoretti, Eliezer M. Van Allen, Toni K. Choueiri, Srinivas R. Viswanathan

**Affiliations:** Department of Medical Oncology, Dana-Farber Cancer Institute, Boston, MA, USA; Broad Institute of MIT and Harvard, Cambridge, MA, USA; Harvard Medical School, Boston, MA, USA; Department of Pathology, Brigham and Women’s Hospital, Boston, MA, USA; Department of Medicine, Massachusetts General Hospital Cancer Center, Boston, MA, USA; Division of Medical Oncology/Hematology, William Osler Health System, Brampton, ON, Canada; Division of Medical Oncology, Tom Baker Cancer Centre, University of Calgary, AB, Canada; Division of Urology, Brigham and Women’s Hospital, Boston, MA, USA; Department of Thoracic/Head and Neck Medical Oncology; Department of Genomic Medicine, MD Anderson Cancer Center, Houston, TX, USA; Department of Urology, Massachusetts General Hospital, Boston, MA, USA; Department of Medicine, Brigham and Women’s Hospital, Boston, MA, USA; Beth Israel Deaconess Medical Center, Boston, MA, USA; Department of Oncologic Pathology, Dana-Farber Cancer Institute, Boston, MA, USA

**Author notes:** Corresponding authors: Toni K. Choueiri, MD, Department of Medical Oncology, Dana-Farber Cancer Institute, 450 Brookline Ave, Boston, Massachusetts, 02215., Srinivas R. Viswanathan, MD, PhD, Department of Medical Oncology, Dana-Farber Cancer Institute, 450 Brookline Ave, Boston, Massachusetts, 02215. Senior authors (Eliezer M. Van Allen, Toni K. Choueiri, Srinivas R. Viswanathan).

**Keywords:** Translocation renal cell carcinoma, genomics, TFE3, TFEB, MITF, NRF2, VEGFR, immune checkpoint inhibition, immunotherapy, oxidative stress

## Abstract

Translocation renal cell carcinoma (tRCC) is an aggressive and poorly-characterized subtype of kidney cancer driven by *MiT/TFE* gene fusions. Here, we define the landmarks of tRCC through an integrative analysis of 152 tRCC patients identified across multiple genomic, clinical trial, and retrospective cohorts. Most tRCCs harbor few somatic alterations apart from *MiT/TFE* fusions and homozygous deletions at chromosome 9p21.3 (19.2% of cases). Transcriptionally, tRCCs display a heightened NRF2-driven antioxidant response that is associated with resistance to many targeted therapies. Consistently, we find that outcomes for tRCC patients treated with vascular endothelial growth factor receptor inhibitors (VEGFR-TKI) are worse than those treated with immune checkpoint inhibition (ICI). Multiparametric immunofluorescence confirmed the presence of CD8 ^+^ tumor-infiltrating T cells compatible with a clinical benefit from ICI and revealed an exhaustion immunophenotype distinct from clear cell RCC. Our findings comprehensively define the clinical and molecular features of tRCC and may inspire new therapeutic hypotheses.

## INTRODUCTION

Translocation renal cell carcinoma (tRCC) is an aggressive subtype of non-clear cell kidney cancer that comprises up to 5% of all RCCs in adults and up to 50% of RCCs in children^1, 2^. Prior case series have suggested that tRCC has a demographic profile that is distinct from more common subtypes of kidney cancer, with a younger age at diagnosis, advanced stage at presentation, and a female predominance^3–,5^. Biologically, tRCCs are driven by activating gene fusions involving transcription factors in the *MiT/TFE* gene family^6–12^. There are currently no molecularly-targeted therapies specific to tRCC and effective treatments for this aggressive cancer remain a major unmet medical need.

A significant barrier to the development of mechanism-inspired therapeutics for tRCC is an incomplete understanding of the molecular landscape and clinical features of the disease. Owing to the rarity of tRCC, prior genomic profiling studies have been limited in scope. While *MiT/TFE* fusions are universal in tRCC, it remains unclear whether there are co-occurring genetic alterations or transcriptional programs that represent additional defining features of the disease^13–15^. Like the molecular landscape, the clinical treatment landscape in tRCC is also largely undefined, with no established standard of care. As a result, tRCC patients are typically treated with therapies originally developed for clear cell RCC (ccRCC)^16^, including vascular endothelial growth factor receptor inhibitors (VEGFR-TKI), multikinase inhibitors (cabozantinib), mTOR inhibitors, or immune checkpoint inhibitors (ICIs). Although some responses to each of these classes of agents have been reported in tRCC, outcomes have been variable between series, and it remains unclear which class(es) of therapeutics are best suited to the biology of tRCC^17–23^.

An intriguing feature of tRCC is that it can exhibit diverse histologic features that may mimic almost all other subtypes of RCC^24, 25^. As a result, tRCC cases have been retrospectively identified within ccRCC and papillary RCC (pRCC) sequencing cohorts^7, 26, 27^. In this study, we leveraged this “histologic overlap” between tRCC and other RCC subtypes to identify tRCC cases from across multiple genomic, clinical trial, and retrospective datasets. We combined these cases with profiling of prospectively identified patients with tRCC to comprehensively characterize the molecular landscape, clinical features, and treatment outcomes for this disease.

## RESULTS

### Identification of tRCC Cases in Large-scale Clinical and Genomic Datasets

To comprehensively characterize both the molecular and clinical features of tRCC, we interrogated RCC cases across multiple large-scale datasets. In a retrospective analysis of metastatic RCC patients from the Dana-Farber/Harvard Cancer Center (Harvard cohort), we identified 734 patients with ccRCC, 97 patients with pRCC, 23 patients with chromophobe RCC (chRCC), and 19 patients with tRCC. tRCC patients were identified on the basis of positive *TFE3* fluorescence *in situ* hybridization (FISH) or strongly positive TFE3 immunohistochemistry with FISH not available. Among this cohort, we observed that tRCC patients had significantly worse outcomes than did patients with the other major histologies of RCC (**Fig. 1a**), a trend that held in an independent metastatic RCC dataset (International Metastatic RCC Database Consortium, IMDC; **Fig. S1a**). Similarly, patients with localized tRCC trended towards the shortest progression-free interval after nephrectomy (**Fig. 1a**). Consistent with smaller case series^3, 5^, we used data from three large independent cohorts (Harvard, IMDC, TCGA) to confirm that tRCCs are female-predominant (**Fig S1b**), present at a younger age (**Fig S1c**), higher stage (**Fig S1d**), and are associated with worse clinical prognostic groups in metastatic disease (**Fig S1e**) as compared with the other major histologies of RCC. Collectively, these data establish tRCC as a disease that predominantly impacts young female patients and is more aggressive than other forms of RCC in both the localized and metastatic settings.

**Fig. 1.**
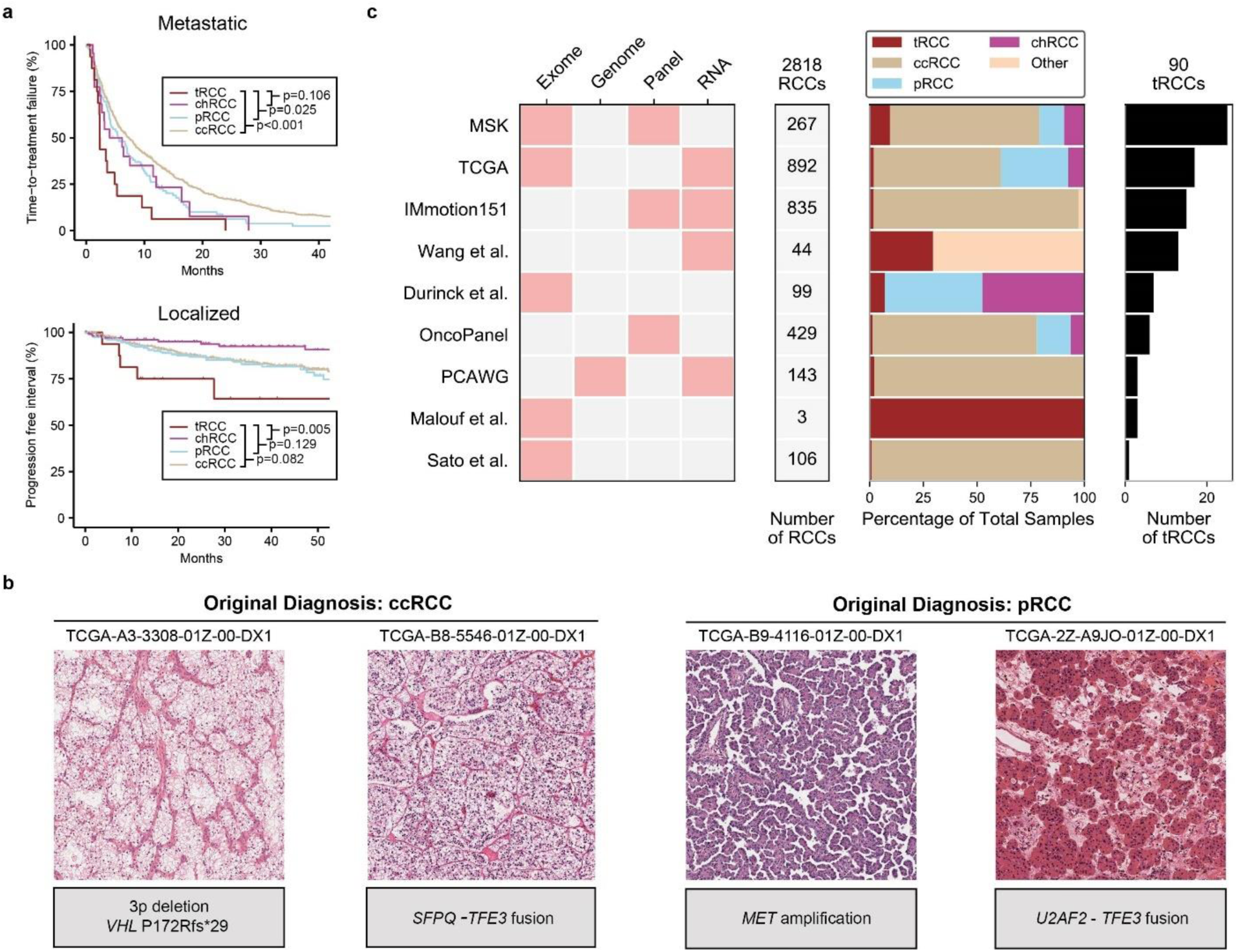
Identification of tRCC cases in multiple clinical and molecular datasets. **a***,Top*, Kaplan-Meier curves for time-to-treatment failure in metastatic ccRCC, pRCC, chrRCC, or tRCC (Harvard cohort). *Bottom*, Kaplan-Meier curves for progression-free interval for localized ccRCC, pRCC, chrRCC, or tRCC (TCGA cohort). P-values were calculated by pairwise log-rank test. **b**, Representative H&E micrographs (x10) of cases originally included in the TCGA ccRCC or pRCC sequencing cohorts. The right case in each pair was subsequently found to have a *TFE3* gene fusion on RNA-Seq. **c,** Aggregation of tRCC cases from across 9 independent NGS datasets. The data type(s) analyzed are indicated for each dataset. tRCC cases were identified based on the presence of a fusion involving an *MiT/TFE* family member (see **Methods**). The number and proportion of tRCC samples as well as number of total RCC samples is indicated for each dataset.

To aggregate tRCC cases for genomic analysis, we leveraged the fact that tRCCs have been reported to share overlapping histologic features with the most frequent histologic subtypes of kidney cancer (ccRCC and pRCC)^28^. As a result, a small number of tRCC cases – harboring defining *MiT/TFE* fusions – have been inadvertently included in several RCC genomic datasets^26, 29–32^. As an example, tRCC cases with histopathologic features indistinguishable from ccRCC and pRCC were included in the Cancer Genome Atlas (TCGA) effort^26, 29^ **(Fig. 1b** and **Supplementary Table 1)**. Building on this observation, we interrogated fusion calls and/or FISH results for 2818 RCCs across 9 independent datasets profiled by DNA sequencing (exome, genome, or panel sequencing) and/or RNA sequencing (**Fig. 1c**). We identified a total of 90 tRCCs with genomic (DNA) or transcriptomic (RNA) profiling data (42 with only genomic data, 16 with only transcriptomic data, 32 with both, **Fig. S1f**).

### Somatic Mutational and Copy Number Alterations in tRCC

We analyzed the 74 tRCC cases on which DNA profiling data were available to elucidate the genomic landscape of tRCC. Among these cases, 36 were profiled via WES, 3 via WGS, and 35 via panel sequencing (**Methods**). tRCC cases showed few mutations overall, with a median (interquartile range) tumor mutational burden of 0.82 (0.43 - 1.28) per megabase (on WES), a rate significantly lower than ccRCC and pRCC and comparable to chRCC (**Fig. S2a),** with similar trends for all (**Fig. S2b**) and frameshift (**Fig. S2c**) indels. Of the most frequently mutated genes in tRCC, none exceeded a frequency of 10% (**Fig. 2a**). These included genes involved in the DNA Damage response (*ATM* (8.1%), *BRCA2* (8.1%), and *WRN* (4.4%)), genes involved in ATP-dependent chromatin remodeling via the SWItch/Sucrose Non-Fermentable (SWI/SNF) complex (ARID1A (5.4%), SMARCA4 (5.4%)), and mutations in *TERT* (6.8%; primarily non-coding mutations in the *TERT* promoter)^33^. Among the 52 cases with gene-level copy number profiling data available, the only recurrent focal alteration in tRCC was homozygous deletion at the *CDKN2A/2B* locus (9p21.3), found in 19.2% of cases. Notably, 50.0% (37/74) of cases in our cohort showed no detectable somatic alterations in either the most frequently mutated tRCC genes or genes that are significantly mutated in clear cell, papillary, or chromophobe RCC **(Fig. 2a)**^27^. Analysis of arm-level copy number alterations among 17 tRCC cases in the TCGA cohort^34^ revealed the most frequent alterations to be hemizygous loss of chromosome 3p (28.6%; though markedly less frequent versus ccRCC 86.8%; p<0.001), chromosome 9p (23.5%), chromosome 18 (29.4%), and chromosome 22q (18.8%), as well as gain of 17q (20.0%) **(Fig. S2e)**. Several of these alterations are defining features of other tumor types of neural/neuroendocrine origin, including monosomy 18 in small intestinal neuroendocrine tumors^35^, 17q gain in neuroblastoma^36^, and 22q loss in pediatric ependymoma^37^.

**Fig. 2.**
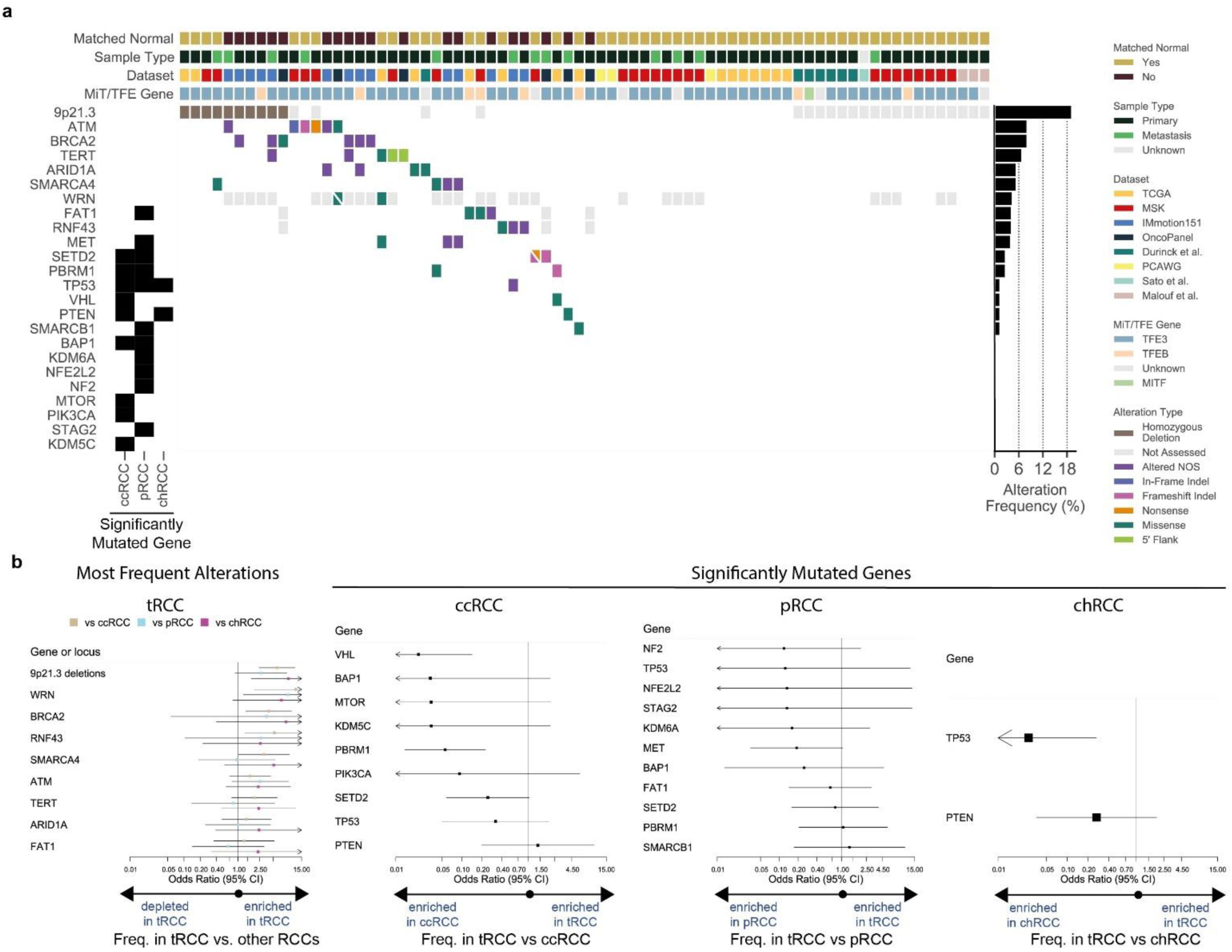
Landscape of genomic alterations in tRCC. **a,** CoMut plot of mutational and copy number alterations in tRCC across all datasets. Genes listed include those found to be most frequently altered in tRCC across all datasets as well as previously reported significantly mutated genes in ccRCC, pRCC, and chrRCC^27^ (indicated in the left track). **b**, Pairwise enrichment analysis for genomic alteration frequencies in tRCC versus other RCC histologies for the indicated genes, presented as the pooled odds ratio and 95% confidence interval from the random-effects meta-analysis in tRCC versus comparator histology. From left to right, genes listed on forest plots indicate: most frequently altered genes in tRCC, significantly mutated genes in ccRCC, significantly mutated genes in pRCC, and significantly mutated genes in chRCC. Pairwise enrichment between tRCC and comparator was calculated individually for each locus or gene within each dataset and pooled estimates across datasets were obtained as detailed in **Methods**.

We next conducted an enrichment analysis of driver gene alteration frequencies between tRCC and other RCC subtypes. We computed pairwise enrichment (tRCC versus ccRCC, pRCC, and chRCC separately) for each locus within each dataset, then used a random-effects meta-analysis to obtain a pooled estimate of gene alteration enrichment or depletion in tRCC versus comparator RCC histologies across datasets (see **Methods**). We found that the genes most frequently altered in tRCC – most notably *CDKN2A/2B* locus (9p21.3) deletions – are highly enriched in tRCC versus other RCC histologies. In contrast, mutations in genes that are significantly mutated in ccRCC, pRCC, and chRCC tended to be depleted in tRCC (**Fig. 2b**). Thus, while tRCCs are genomically quiet overall (with a lower mutational and copy number alteration burden than other RCC histologies), a subset harbor recurrent alterations -- distinct in profile from those seen in other RCCs -- that may cooperate with the *MiT*/*TFE* fusion to drive cancer.

### Structure of *MiT/TFE* fusions in tRCC

We next turned our attention to further analysis of the *MiT/TFE* fusion, the defining genetic lesion in tRCC. Across the combined tRCC cohort, we found that the vast majority of cases (78 cases; 88.6%) harbored *TFE3* fusions, while the remainder harbored *TFEB* (8 cases; 9.1%) or *MITF* (2 cases; 2.3%) fusions (**Fig. 3a**). Seventeen different *MiT/TFE* fusion partners were observed across the cohort and the spectrum of fusion partners was largely distinct between *TFE3*, *TFEB*, and *MITF* (**Fig. S3a**). The most common *TFE3* fusion partners were *ASPSCR1, SFPQ, PRCC,* and *NONO.* Interestingly several chromosomes harbored multiple potential *MiT/TFE* fusion partners (chr1, chr17, chrX) (**Fig. 3b**). *MiT/TFE* fusion partners showed an enrichment for ontology terms involving RNA processing and RNA splicing, and this was driven predominantly by *TFE3* fusion partners (**Fig. 3c** and **Fig. S3b-c**). Analysis of fusion breakpoints revealed that all fusions preserved the C-terminal helix-loop-helix/leucine zipper domain (HLH-LZ) of the MiT/TFE transcription factor, the region of the protein critical for dimerization and DNA binding^38^; the activation domain was variably preserved in the fusion product (**Fig. 3d** and **Supplementary Table 2)**. Interestingly, large N-terminal portions of most TFE3 fusion partners were included in the fusion, including, domains with RNA-binding potential in cases where the fusion partner was an RNA binding protein. In contrast, TFEB and MITF fusion partners tended to preserve less of the N-terminal fusion partner in the fusion product (**Fig. 3e**). Overall, our results point to a coherent logic to the structure of *MiT/TFE* fusions despite great diversity in fusion partners and breakpoints.

**Fig. 3.**
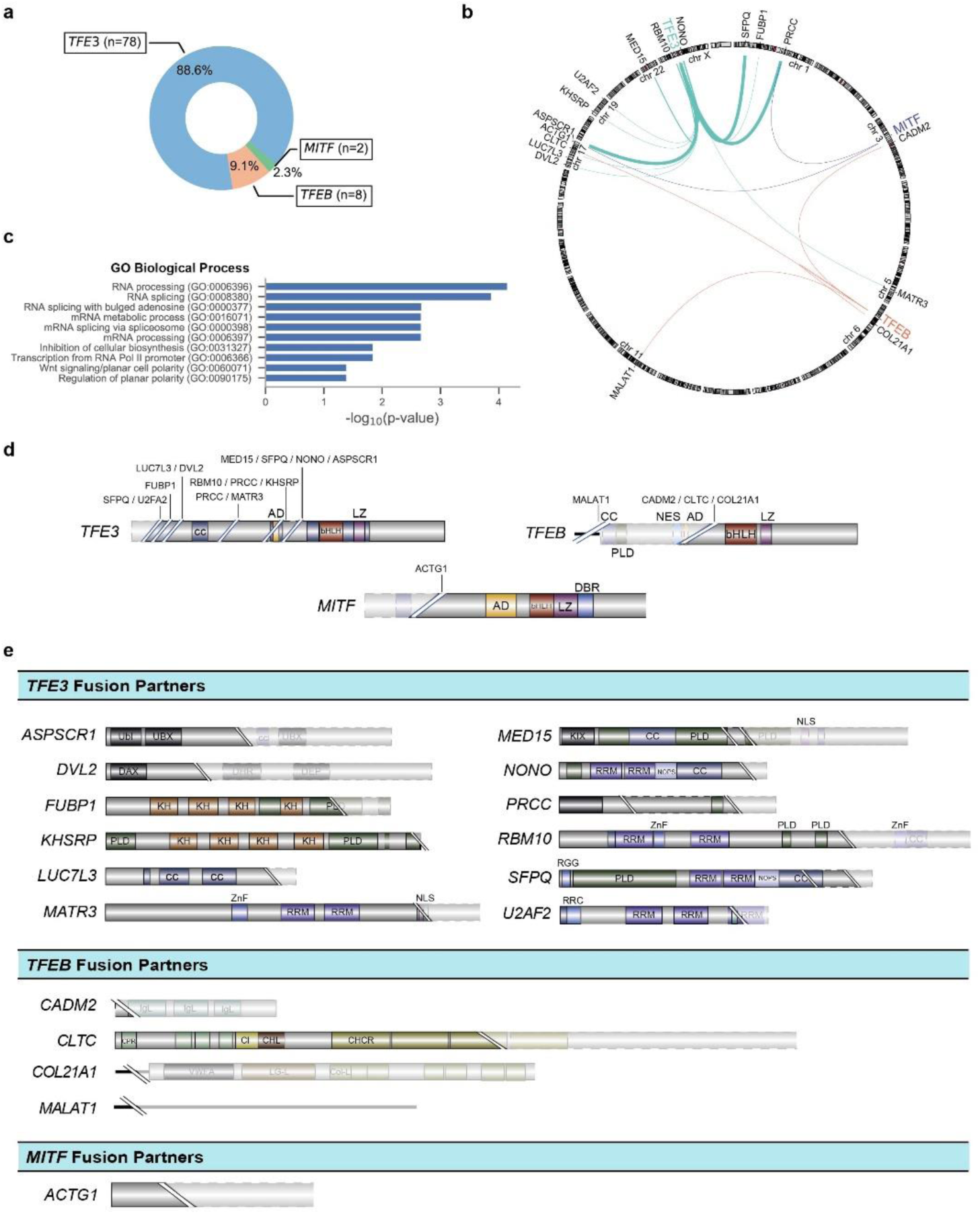
Structure of *MiT/TFE* fusions in tRCC. **a**, Number and percentage of tRCC cases displaying gene fusions involving *TFE3*, *TFEB*, or *MITF* across all datasets analyzed. **b**, Genomic location of *MiT/TFE* fusion partners. Stroke thickness is proportional to the number of times a given gene was observed to be an *MiT/TFE* fusion partner across all datasets analyzed. **c**, Gene ontology terms (GO Biological Process) enriched amongst *MiT/TFE* fusion partners. **d**, Breakpoints observed within *TFE3*, *TFEB, or MITF* across all samples analyzed. Solid portion represents the portion of the *MiT/TFE* gene retained within the oncogenic fusion product. Fusion partner genes observed to join at a given breakpoint are listed. Functional domains within each MiT/TFE gene are indicated (legend in **Supplementary Table 2**). **e**, Breakpoints observed within *MiT/TFE* partner genes. Solid portion represents the portion of each partner gene retained within the oncogenic fusion product. Genes are grouped by whether they were observed to fuse with *TFE3* (top), *TFEB* (middle), or *MITF* (bottom). Functional domains within each *MiT/TFE* partner gene are indicated (legend in **Supplementary Table 2**).

### Distinctive transcriptional features of tRCC

Given our observation that most tRCCs harbor few genomic alterations aside from the *MiT/TFE* fusion, we next sought to determine whether the transcriptional program of tRCC is largely driven by the fusion. We ectopically expressed either wild type (WT) *TFE3* or four of the most common *TFE3* fusions (*ASPSCR1-TFE3*, *NONO-TFE3*, *PRCC-TFE3*, *SFPQ-TFE3*) in 293T cells and performed RNA-Seq (**Fig. 4a** and **Supplementary Table 3**). We derived a 139-gene transcriptional signature based on genes differentially expressed upon *TFE3* fusion, but not WT *TFE3*, expression (**Fig. S4a, Supplementary Table 4** and **Methods**). Subsequently, we performed unsupervised hierarchical clustering using this fusion-specific signature. We observed that tRCC samples clustered tightly together across four independent datasets^30, 39–41^ (**Fig. 4b** and **Fig. S4b**). Clustering based on our fusion-derived signature resulted in superior grouping of tRCCs than did clustering based on the 1000 most variable genes in each dataset (**Fig. S4c**). We then performed differential expression analysis to identify a consensus set of genes overexpressed in tRCC as compared with all comparator tumor types. In each dataset, we performed pairwise comparisons between tRCC and each comparator tumor type to identify genes selectively overexpressed in tRCC (q-value <0.05; **Fig. S4d-e**). We identified a consensus list of 76 genes that were selectively overexpressed in tRCC (q-value <0.05) in 9/13 or more pairwise comparisons (**Fig. 4c** and **Fig. S4e**). Notably, several of these have been previously annotated as MITF target genes^42, 43^ on the basis of prior ChIP-Seq studies and include genes involved in neuronal development (*SNCB, TRIM67*, *IRX6*)^44–46^, ion flux and the antioxidant stress response (*SQSTM1*, *TMEM64, SLC39A1*)^46–48^, and lysosomal function/mTORC1 signaling (*RAB7A*, *RHEB*, *RRAGC*, *ATP6V1C1*)^49–51^. We performed gene set enrichment analysis (GSEA)^52^ using hallmark gene sets^53^ to identify pathways selectively activated in tRCC. This revealed a strong enrichment for gene sets pertaining to reactive oxidative species (ROS) sensing and the response to oxidative stress and xenobiotics (top tRCC-enriched gene sets shown in **Fig. 4d).** In sum, the transcriptional program of tRCC appeared to be driven by the MiT/TFE fusion and resulted in overexpression of genes implicated in mTORC1 signaling, antioxidant stress response, ROS sensing, and the response to oxidative stress and xenobiotics.

**Fig. 4.**
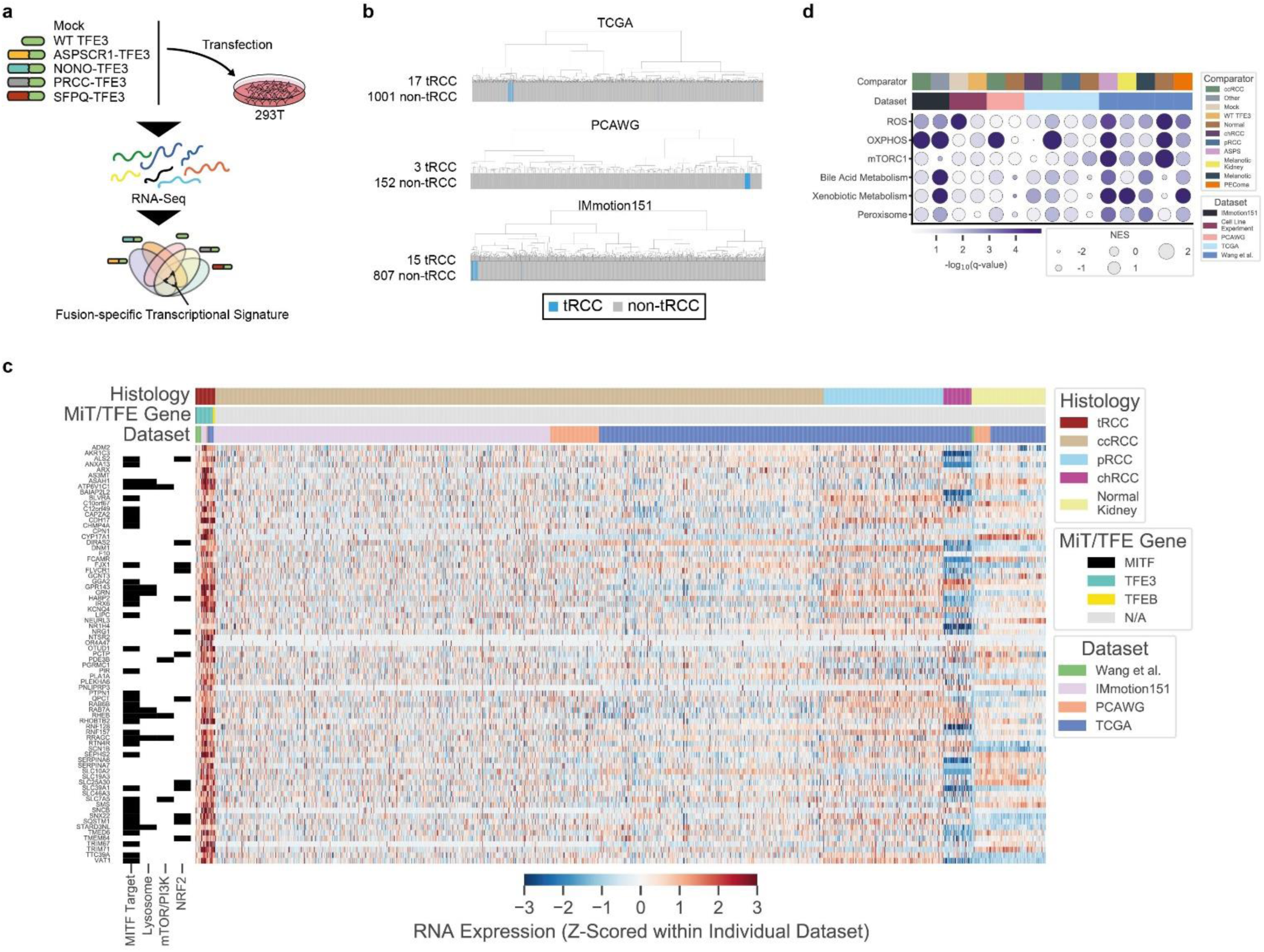
Distinctive transcriptional features of tRCC. **a,** Schematic of *in vitro* experiment used to derive *TFE3*-fusion-specific transcriptional signature. **b**, Transcriptome sequencing data from three independent datasets (TCGA, PCAWG, IMmotion151) were subjected to unsupervised hierarchical clustering using the fusion-specific signature derived in (a). Blue bars indicate *MiT/TFE*-fusion-positive cases within each dataset. Gray bars indicate other RCC histologic subtypes or normal kidney. **c**, Heatmap of genes overexpressed in tRCC as compared with other RCC subtypes or normal kidney, across all datasets (see **Fig.S4**). Membership of genes in key pathways related to tRCC pathogenesis is indicated in the track at left. **d**, Gene set enrichment analysis showing top enriched Hallmark pathways in tRCC samples versus comparators across all datasets analyzed. Dataset and pairwise comparison across which the GSEA was performed is indicated in the track at the top of each column. Dot size is proportional to normalized enrichment score (NES) in tRCC versus comparator; dot color reflects - log10(q-value) for the enrichment.

### An antioxidant response signature associated with resistance to targeted therapies in tRCC

The transcription factor NRF2 (nuclear factor erythroid-derived-2-like 2, *NFE2L2*) is a master regulator of the cellular antioxidant response and controls the expression of genes involved in the response to xenobiotics and oxidative stress^54^. Notably, activation of the NRF2 pathway has been reported in certain subsets of RCC via diverse mechanisms that include somatic alteration or hypermethylation of NRF2 pathway members^55^ and the production of oncometabolites that modify and inhibit KEAP1, a negative regulator of NRF2^7, 10, 27^. Given evidence of activated ROS-sensing in tRCC (**Fig. 4c-d**), we derived an NRF2 activity score using single sample GSEA (ssGSEA)^56^ (based on a 55-gene NRF2 signature^57^) across all RCC samples with available transcriptome profiling data (46 total tRCC samples across 4 datasets; NRF2 activity calculated and Z-scored separately within each individual dataset). We observed that NRF2 activity was universally high amongst tRCC samples as compared with other RCC types and normal kidney tissue (**Fig. 5a**).

**Fig. 5.**
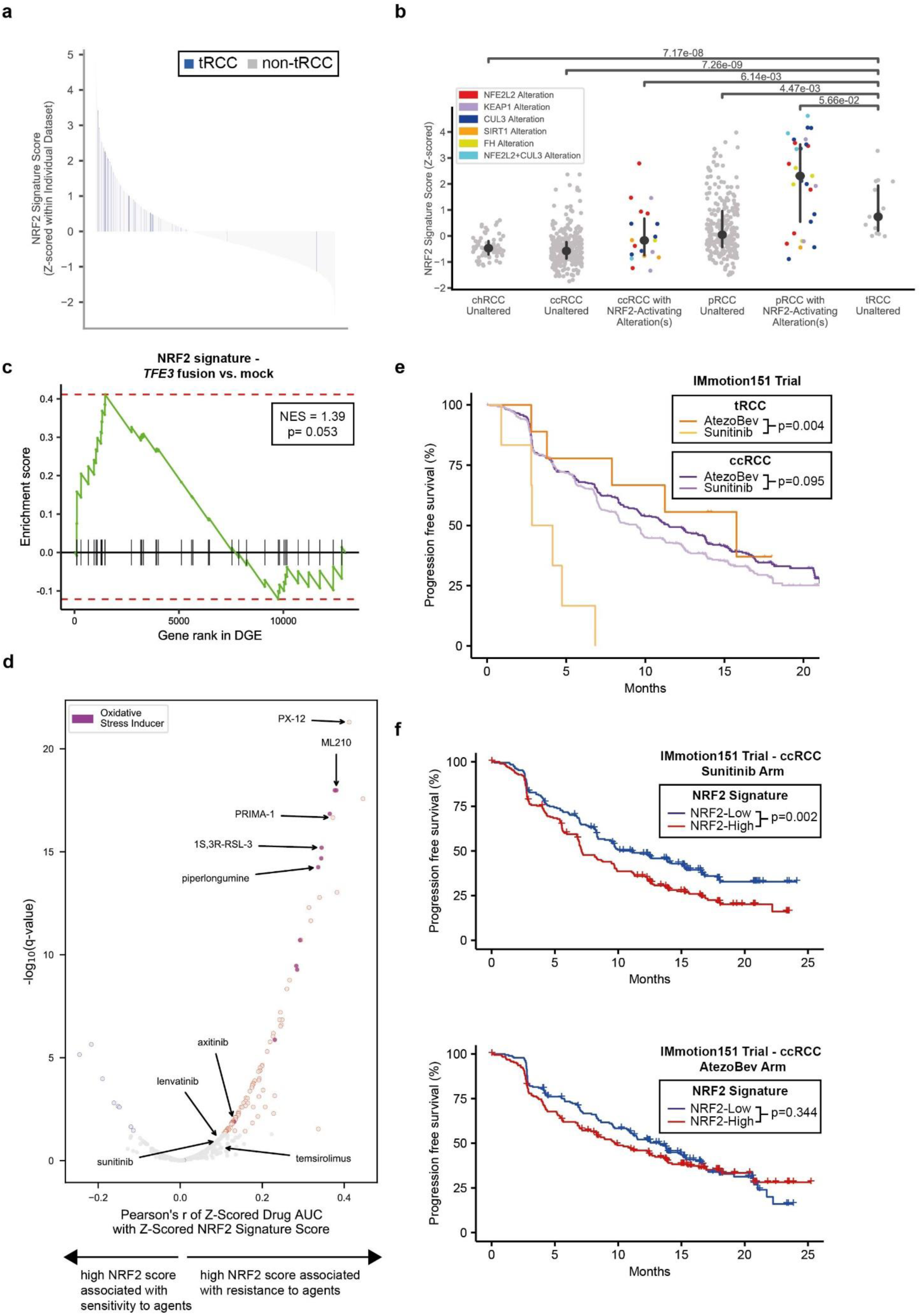
tRCC displays activated NRF2 pathway signaling and a relative resistance to targeted therapies. **a,** Waterfall plot showing NRF2 signature score for all RCC samples across all datasets analyzed. tRCC samples are depicted in blue (n=46); other samples (ccRCC, pRCC, chRCC, normal kidney, or other tumors) are shown in gray (n=1999). **b**, NRF2 signature score for TCGA RCC samples of the indicated histologies. For each histology, samples with somatic alterations in the NRF2 pathway are shown separately. No chRCC or tRCC samples displayed somatic alterations in the NRF2 pathway. **c**, Gene set enrichment analysis showing enrichment of NRF2 gene signature in 293T cells expressing *TFE3* fusions versus mock (untransfected) control condition. **d**, Volcano plot showing correlation of NRF2 signature score with drug sensitivity in the Broad Institute Cancer Therapeutics Response Portal dataset^133^. A high NRF2 signature score is significantly associated with resistance to the agents shown in red. Agents annotated to act through the induction of oxidative stress or ferroptosis are colored in purple. Selected targeted agents used in the treatment of kidney cancer are labeled. **e**, Progression-free survival curves for tRCC (dark and light orange) or ccRCC (dark and light purple) patients treated with either atezolizumab and bevacizumab (AtezoBev) or sunitinib in the randomized Phase III IMmotion151 trial. **f**, Progression-free survival curves for ccRCC patients with high (red) or low (blue) NRF2 signature score treated with either sunitinib (top) atezolizumab + bevacizumab (bottom) on the IMmotion151 trial. For **e-f**, NRF2 signature score was dichotomized at the median in each arm.

We next investigated whether high NRF2 activity in tRCC was attributable to somatic alterations in this pathway. We observed that somatic alterations in the NRF2 pathway (most commonly *KEAP1* or *NFE2L2* alteration) were associated with an increased NRF2 activity score in ccRCC and pRCC, as was a CpG island methylator phenotype (CIMP), consistent with prior reports (**Fig. 5b** and **Fig. S5a**)^27^. Interestingly, however, tRCC samples showed uniformly elevated NRF2 activity, comparable to ccRCC/pRCC samples with somatic alterations in the NRF2 pathway (**Fig. 5b**), despite having no detectable NRF2 pathway alterations. The expression of strong oncogenes has been linked to NRF2 pathway activation^58^ and our transcriptomic analyses revealed overlapping targets between NRF2 and MITF (**Fig. 4c**, hypergeometric one-tailed p-value< 0.001). Consistently, we observed that the NRF2 gene signature was enriched upon ectopic expression of all *TFE3* fusions in 293T cells as compared to the mock treatment condition, suggesting that expression of the *TFE3* fusion may be directly linked to activation of the NRF2 pathway (**Fig. 5c**).

Activation of the NRF2 pathway has been associated with resistance to a number of ROS-producing drugs, including inducers of ferroptosis, a regulated form of iron-dependent oxidative cell death^57, 59, 60^. We calculated a correlation between NRF2 activity score and drug sensitivity across 593 cell lines and 481 compounds assayed in the Cancer Therapeutics Response Portal^61^. Strikingly, high NRF2 activity was associated with relative resistance to almost all agents assayed, including several targeted therapies used in the treatment of RCC (e.g. sunitinib, axitinib, lenvatinib, temsirolimus), and most notably, to multiple compounds known to induce electrophilic stress and oxidative cell death (e.g. PRIMA-1, PX-12, piperlongumine, ML-210, RSL-3) (**Fig. 5d**)^62^. In order to uncover potential vulnerabilities of this otherwise drug-resistant state, we next surveyed pooled genetic (shRNA and CRISPR) screening data generated as part of the Cancer Dependency Map effort^63, 64^ . In both the CRISPR and shRNA datasets, we found that the outlier dependency of NRF2-high cells is *NFE2L2* itself (**Fig. S5b**). Although tRCC cell lines are not currently included among those assayed in the Cancer Dependency Map effort, we separately validated that three tRCC cell lines all demonstrated variable levels of dependency on *NFE2L2* knockdown, consistent with the notion that direct inhibition of NRF2 is a vulnerability of the NRF2-high state observed in tRCC (**Fig. S5c**).

Next, to determine whether elevated NRF2 activity might be associated with resistance to targeted therapies in patients, we evaluated molecular data from the IMmotion151 trial (NCT02420821), a Phase III trial of 915 RCC patients with clear cell or sarcomatoid histology who were randomized to either sunitinib (multitargeted kinase inhibitor against VEGFRs and PDGFRs) or the combination of atezolizumab (monoclonal antibody targeting PD-L1) and bevacizumab (monoclonal antibody targeted VEGF-A)^65^. RNA-Seq performed on tumor biopsies from patients enrolled on this trial revealed 15 patients with *TFEB/TFE3* translocations among 822 with available RNA-seq data (**Fig. 1c**), of which 6 were treated on the sunitinib arm and 9 were treated on the atezolizumab + bevacizumab (AtezoBev) arm^30^. While AtezoBev showed a modest benefit over sunitinib in progression-free survival (PFS) in the overall study and amongst ccRCC patients, we observed that tRCC patients receiving sunitinib did dramatically worse than those receiving AtezoBev (median PFS 3.5 months with sunitinib vs. 15.8 months with AtezoBev; log-rank p= 0.004). Consistent with this observation, the extent of benefit derived from AtezoBev as compared with sunitinib, in patients with tRCC vs. ccRCC, was significantly greater (histology-by-treatment arm interaction Cox p-value=0.008) (**Fig. 5e**). When ccRCC patients treated with sunitinib were dichotomized based on NRF2 activity score, those with high-NRF2 scores had shorter PFS compared to with low-NRF2 scores (median PFS 7.1 months for high-NRF2 vs. 11.1 months for low-NRF2; log-rank p=0.002). In contrast, NRF2 activity score was not associated with a significant difference in outcome in ccRCC patients treated on the AtezoBev arm (**Fig. 5f**). In the CheckMate cohort including 311 patients with ccRCC with available RNA-seq data (pooled analysis of the CheckMate 009 [NCT01358721], 010 [NCT01354431], and 025 [NCT01668784] clinical trials)^66^, a similar signal was observed whereby ccRCC patients with a high NRF2 activity score experienced shorter PFS than did those with a low NRF2 activity score (**Fig. S5d**), on the everolimus arm (median PFS 9.7 months for high-NRF2 vs. 14.3 months for low-NRF2; log-rank p= 0.031), but not the nivolumab arm^67^. Together, these results indicate that high NRF2 activity – a defining feature of tRCC – is associated with resistance to targeted agents used in the treatment of RCC, but may not preclude responses to ICI.

### Response to immune checkpoint inhibition in tRCC

We sought to further explore the possibility that tRCC may be responsive to ICI. Analysis of responses from the IMmotion151 study showed that tRCC patients derived significantly greater clinical benefit (CB) on AtezoBev than on sunitinib (77.8% with AtezoBev vs. 16.7% with sunitinib; Fisher p-value= 0.041). However, tRCC patients tended to not derive clinical benefit (no clinical benefit; NCB) from sunitinib as compared with AtezoBev (11.1% with AtezoBev vs. 50.0% with sunitinib; Fisher p-value= 0.235). In contrast, ccRCC patients tended to have similar CB (65.1% with AtezoBev vs. 64.0% with sunitinib; Fisher p-value= 0.767) and NCB (15.6% with AtezoBev vs. 16.0% with sunitinib; Fisher p-value= 0.923) rates whether they received AtezoBev or sunitinib (**Fig. 6a**).

**Fig. 6.**
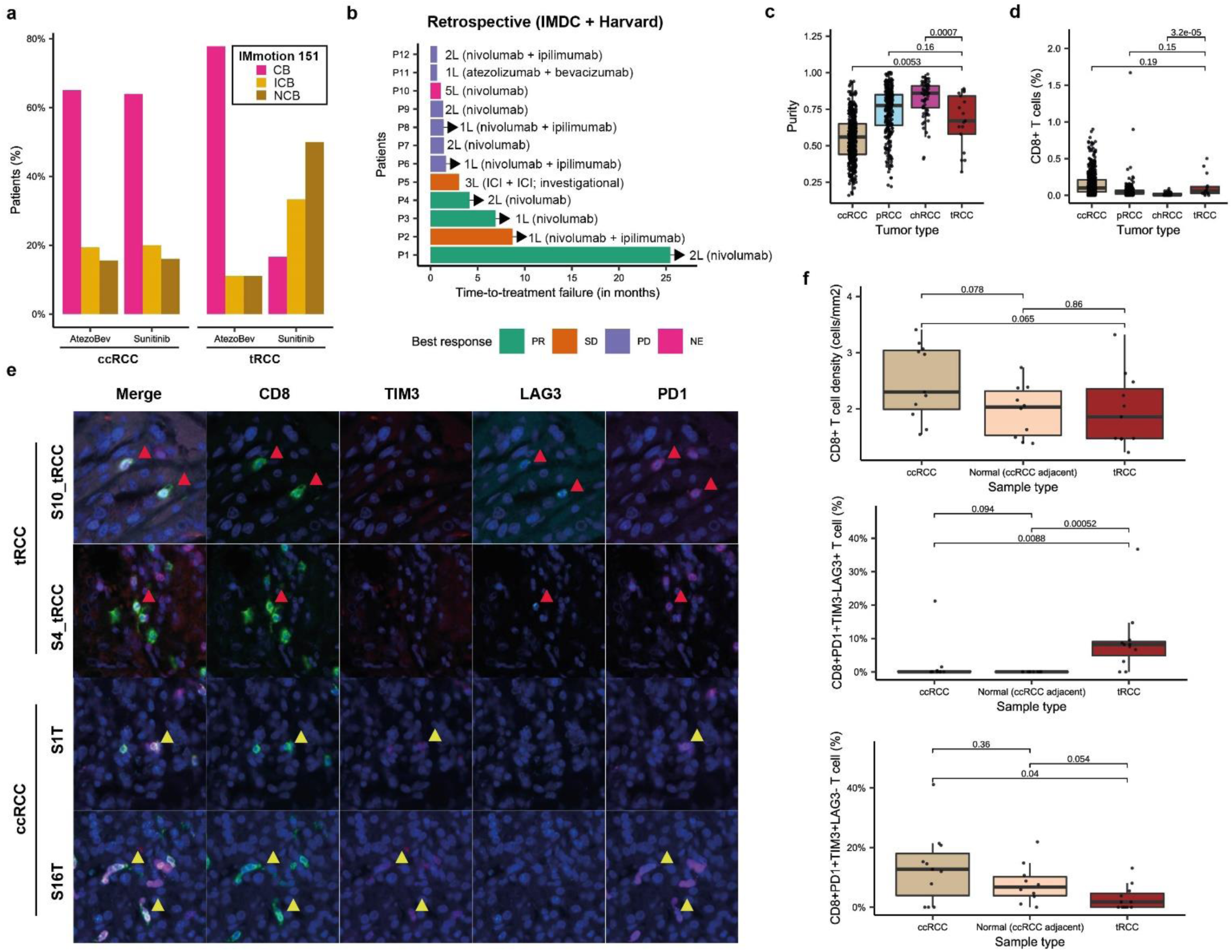
Immunogenomic features of tRCC associated with responses to immune checkpoint inhibition. **a,** Percentage of tRCC patients showing clinical benefit (CB), intermediate clinical benefit (ICB), or no clinical benefit (NCB) to either AtezoBev or sunitinib on the IMmotion151 trial. **b**, Swimmer plot showing response types and response times to immune checkpoint inhibitor-based regimens in tRCC patients in the combined IMDC and Harvard retrospective cohort. Line (L) in which ICI was received as well as specific ICI regimen is indicated to the right of each patient. **c**, Sample purity in tRCC, ccRCC, chRCC, and pRCC in the TCGA cohort. **d**, CD8^+^ T cell infiltration imputed from gene expression (CIBERSORTx) in tRCC, ccRCC, chRCC, and pRCC in the TCGA cohort. **e**, Multiparametric immunofluorescence for CD8, TIM3, LAG3, and PD1 in representative tRCC cases (top two rows) and ccRCC cases (bottom two rows). Red arrows indicate CD8^+^PD1^+^LAG3^+^TIM3^-^ tumor-infiltrating T cells in tRCC cases. Yellow arrows indicate CD8^+^PD1^+^LAG3^-^TIM3^+^ tumor-infiltrating T cells in ccRCC cases. **f**, Quantification of CD8^+^ T-cell density (top), percentage of CD8^+^PD1^+^TIM3^-^LAG3^+^ T cells (middle), and percentage of CD8^+^PD1^+^TIM3^+^LAG3^-^ T cells (bottom) in tRCC (n= 11), ccRCC (n= 11), and adjacent normal tissue (from ccRCC cases, n= 10).

In a combined analysis of the IMDC and Harvard datasets, we identified 12 metastatic tRCC patients who had received ICI in any line of therapy as well as 10 tRCCs that had been treated by TKIs (n= 8 sunitinib; n= 2 pazopanib). Among this cohort, 5 achieved either partial response (n= 3) or stable disease (n= 2) on an ICI-containing regimen, with several ongoing responses (**Fig. 6b** and **Fig. S6a-b**). Overall, in this retrospective combined cohort of tRCC patients, the response rate (25.0% with ICI and 0% with TKI; Fisher p-value= 0.220) and overall survival (OS; median OS 62.4 months with ICI and median OS 10.3 months with TKI; log-rank p-value= 0.267) tended to be increased on ICI-based regimens compared to TKIs (**Fig. S6c-d**), corroborating the result that tRCC patients may derive greater benefit from ICI-based therapies than VEGF-targeted therapies.

We next examined whether immunogenomic features of tRCC could explain responses to ICI in this RCC subtype, despite a low burden of mutations and CNAs (**Fig. 2a** and **Fig. S2**). In the TCGA cohort, tumor purity (which is inversely correlated to immune cell infiltration), was lower in tRCC than chRCC (a classically ICI-resistant subtype^68, 69^) (Wilcoxon p-value< 0.001), similar to pRCC (Wilcoxon p-value= 0.160), and higher than ccRCC (Wilcoxon p-value= 0.005) (**Fig. 6c**). Consistently, immune deconvolution analyses (CIBERSORTx^70^) showed that the inferred percentage of cluster of differentiation 8 (CD8)^+^ T cells was higher in tRCC than in chRCC (Wilcoxon p-value< 0.001), and comparable to that seen in ccRCC (Wilcoxon p-value= 0.190) and pRCC (Wilcoxon p-value= 0.150) (**Fig. 6d**). Additionally, PD-L1 protein levels on tumor-infiltrating immune cells, as assessed by IHC, in patients on the IMmotion151 trial, were comparable between tRCC and ccRCC patients (41.8% with ccRCC vs. 33.3% with tRCC; Fisher p-value= 0.604) (**Fig. S6e)**.

Finally, we sought to more carefully characterize the CD8^+^ tumor-infiltrating T cells in tRCC via multiparametric immunofluorescence^71, 72^. We examined 11 ccRCC cases (including 10 with adjacent normal tissue) and 11 tRCC cases for T cells expressing CD8 or the immune checkpoint markers PD1, T-cell immunoglobulin and mucin-domain containing-3 (TIM3), and lymphocyte activation protein-3 (LAG3). While the overall CD8^+^ T cell density tended to be lower in tRCC samples than in ccRCC samples (Wilcoxon p-value = 0.065) (**Fig. 6e-f**), the percentage of CD8^+^PD1^+^TIM3^-^LAG3^-^ cells (the subset predictive of a response to PD1/PD-L1-based ICI^71, 72^) was not significantly different between tRCC and ccRCC (**Fig. S6f**). Moreover, the profile of immune checkpoint markers differed significantly between ccRCC and tRCC; tRCC cases displayed a higher percentage of CD8^+^PD1^+^TIM3^-^LAG3^+^ T cells (Wilcoxon p-value = 0.009) whereas ccRCC cases displayed a higher percentage of CD8^+^PD1^+^TIM3^+^LAG3^-^ T cells (Wilcoxon p-value = 0.040). Altogether, our results are consistent with the notion that tRCCs may benefit from ICI as a result of a permissive immune microenvironment characterized by a tumor-infiltrating T cell profile distinct from that observed in ccRCC.

## DISCUSSION

We performed a comprehensive and multicenter characterization of the molecular and clinical features of 152 tRCCs. While prior studies have identified some genomic and transcriptional features of tRCC, the broader extensibility of these findings, their clinical actionability, as well as an understanding of how they compare to other subtypes of RCC have remained unclear^13–15^. Our integrative analysis spans genomic and transcriptomic data, immunophenotypic analysis, functional validation, and clinical outcome data from both retrospective cohorts and randomized clinical trials. From these efforts, an increasingly well-defined landscape of tRCC emerges.

The defining – and often singular – genomic alteration in tRCC is the *MiT/TFE* fusion. Our results show that *TFE3* is by far the most frequently involved *MiT/TFE* gene. While there exists a great diversity of *MiT/TFE* fusion partners, these partners are highly enriched on certain chromosomes (chr1, chr17, chrX), raising intriguing questions about whether patterns of spatial genome organization underlie these recurrent translocations^73–75^. Moreover, our analysis of breakpoint locations across fusions highlights that the vast majority of *TFE3* fusions arise via in-frame events that preserve functional domains from both *TFE3* and its partner protein (most of which are RNA binding proteins); this opens the possibility that *TFE3* fusion partners may confer neomorphic activity to the fusion product. In contrast, much smaller regions of *TFEB* and *MITF* partner genes appear to be involved in the fusion product. Whether differences in fusion structure translate to histologic and/or phenotypic differences between *TFE3*-, *TFEB*-, and *MITF*-translocation RCC warrants further investigation^1, 76, 77^.

Overall, tRCCs are genomically quiet tumors with a low mutational and copy number alteration burden, a reduced frequency of alterations in genes known to be significantly mutated in other RCC subtypes, and few recurrent alterations aside from the *MiT/TFE* fusion. A notable exception is homozygous loss at chromosome 9p21.3, which harbors the *CDKN2A/2B* genes, and is found in 19.2% of tRCC cases. Loss of CDKN2 proteins may be associated with high CDK4/6 activity and may sensitize to CDK4/6 inhibitors^78^. Co-deletion of *MTAP*, which is located in close proximity to *CDKN2A*, may sensitize to PRMT5 inhibitors^79, 80^. Mutations in *TERT* (primarily in the promoter region) were also found in 6.8% of cases. Notably, both *CDKN2A/B* loss and *TERT* promoter mutations are defining genetic features of malignant melanoma, a cancer type driven by activated MITF signaling^33, 81–83^. Less frequent alterations in the cohort included multiple genes involved in the DNA damage response (*ATM*, *BRCA2*, *WRN*), though the lack of specific variant information, the absence of matched normal-based filtering of mutation calls for some samples, and low alteration frequency preclude drawing strong conclusions about this class of mutations.

We identified a heightened response to oxidative stress as a transcriptional hallmark of tRCC. Activated NRF2 signaling has been linked to oncogenesis and resistance to chemotherapies in various contexts^84^. Prior studies have indicated that small subsets of both ccRCC and pRCC display heightened NRF2 signaling, generally linked to somatic alterations or DNA methylation in the NRF2 pathway^7, 27, 85^. Interestingly, our results suggest that NRF2 signaling is uniformly activated in tRCC in the absence of detectable somatic alterations in the NRF2 pathway. Notably, multiple NRF2 target genes are also annotated as MiT/TFE targets (**Fig. 4c**), suggesting a direct link between MiT/TFE fusions and the NRF2 pathway in tRCC. Our results may explain why tRCCs (and ccRCCs with elevated NRF2 signaling) display worse outcomes with sunitinib than with ICI in clinical datasets, and are consistent with *in vitro* data suggesting that NRF2 confers resistance to sunitinib and other TKIs^55, 86, 87^. Whether this signal holds for extended spectrum kinase inhibitors such as cabozantinib and lenvatinib remains to be determined, as patients receiving these therapies were not represented in our retrospective cohort. We validate that *NFE2L2* represents a clear genetic dependency of the NRF2-high state, and suggest that specific NRF2 pathway inhibitors, if developed, may be effective in tRCC^54, 88^.

Responses to ICI in tRCC are notable given the apparent lack of potential sources of tumor-associated antigens (i.e. low burden of mutations and indels). Our immune deconvolution analyses and immunofluorescence studies both support the notion that tRCCs do contain an appreciable density of tumor-infiltrating CD8^+^ T cells. The tumor neoantigens recruiting T cells in tRCC may be derived from the fusion junction, as has also been reported for other fusion-driven malignancies^15, 89^. Interestingly, there is no significant difference in the percentage of CD8^+^PD1^+^TIM3^-^LAG3^-^ T cells – the activated non-exhausted T-cell subset that is implicated in an effective antitumor response – between ccRCC (a classically ICI-responsive tumor) and tRCC^90–92^. The immunophenotype of exhausted T cells also appears to differ between ccRCC and tRCC: CD8^+^PD1^+^TIM3^-^LAG3^+^ T cells are predominant in tRCC while CD8^+^PD1^+^TIM3^+^LAG3^-^ T cells are predominant in ccRCC. Both TIM3 and LAG3 have been proposed as immune checkpoints that can be targeted in combination with PD-1/PD-L1. Notably, several trials combining LAG3 blockade with PD1 blockade are currently underway (and include patients with RCC)^90^ and this combination has recently shown to have efficacy in patients with previously untreated metastatic melanoma^93^. Our immunophenotypic data provide rationale for the development of this therapeutic combination in tRCC. Our findings are also consistent with those of a prior study that showed, using a lung adenocarcinoma mouse model, that activated NRF2 and PI3K/mTOR signaling can lead to changes in the immune microenvironment that are permissive to ICI response^94^. In tRCC, our results suggest that both the PI3K/AKT/mTOR pathway and NRF2 may be activated downstream of MiT/TFE fusions (**Fig. 4c**)^21^.

Our study does have several limitations. First, the cohort is heterogeneous in terms of stage of disease (localized and metastatic), sequencing platform used, and data types available for analysis. While the heterogeneity of the cohort is inevitable given the rarity of the disease, the analysis methods we apply account for dataset-specific biases (**Methods**) and the scale of this study has enabled us to make multiple novel insights. Second, tRCCs are themselves a heterogeneous group of tumors with respect to fusion partners, biology, and prognosis^95^. Larger studies or more homogeneous cohorts comprised of prospectively collected samples will be required to draw strong conclusions about how the specific *MiT/TFE* gene or its fusion partner influence disease biology. Third, some of our clinical data are retrospective, which has inherent limitations. Nonetheless, we suggest that the signals observed from misclassified tRCC patients enrolled on randomized clinical trials for ccRCC, and the corroboration of these signals by translational and retrospective clinical data, may have important implications for the treatment of tRCC.

Altogether, we demonstrate the power of integrative clinico-genomic analysis to illuminate the molecular underpinnings and clinical features of tRCC. Our work inspires multiple hypotheses that can be pursued in future studies to further dissect the biology of this rare cancer. These data also lay the framework for the development and testing of mechanism-driven therapeutic regimens in tRCC.

## METHODS

### Clinical tRCC cohorts

The comparison of baseline characteristics and clinical outcomes was done using data from patients included in two retrospective cohorts of consecutive patients: (1) Harvard cohort (n= 734 ccRCC, n= 97 pRCC, n= 23 chRCC, n= 19 tRCC), a retrospective cohort from the Dana-Farber/Harvard Cancer Center including patients from Dana-Farber Cancer Institute, Beth Israel Deaconess Medical Center, and Massachusetts General Hospital and (2) IMDC cohort (n= 6107 ccRCC, n= 396 pRCC, n= 107 chRCC, n= 40 tRCC): a retrospective multi-center cohort of metastatic RCC that includes more than 40 international cancer centers and more than 10,000 patients with metastatic RCC^96^. All patients consented to an institutional review board (IRB) approved protocol to have their clinical data retrospectively collected for research purposes and the analysis was performed under a secondary use protocol, approved by the Dana-Farber Cancer Institute IRB. For the Harvard cohort, tRCC patients were defined as: (1) positive *TFE3* FISH test or (2) positive TFE3 test by IHC along with a strongly suggestive clinico-pathologic history and no FISH testing results available (missing). For the IMDC cohort, patients were included as tRCCs if they (1) had a positive *TFE3* FISH test, (2) had a positive TFE3 IHC test and suggestive clinico-pathologic history and no FISH testing data available (missing), or (3) no *TFE3* FISH or TFE3 IHC test results available but suggestive clinico-pathologic history. Clinico-pathologic diagnoses were used to define comparator RCC histologies (ccRCC, pRCC, and chRCC). For the IMDC cohort, comparator histologies (controls) were only used from clinical sites that contributed tRCC cases.

### Genomic tRCC cohorts

For genomic datasets, tRCCs were identified based on RNA-seq-based fusion calls, a positive *TFE3* FISH test, or DNA-based fusion calls derived from panel data (MSK-IMPACT or OncoPanel). Clinico-pathologic diagnoses were used to define the cases of other RCC histologies (ccRCC, pRCC, chRCC, normal kidney, or other). Data for the Memorial-Sloan Kettering (MSK) cohort was obtained from the study by Marcon et al.^15^ and Zehir et al.^97^. Fusion calls for the TCGA cohort were obtained from the study by Gao et al.^29^, clinico-pathologic data was obtained from Genomic Data Commons (https://gdc.cancer.gov/about-data/publications/pancanatlas), and the pathology slides used in **Fig. 1b** were obtained from https://portal.gdc.cancer.gov/. Data for the PCAWG^98^ cohort were obtained from the ICGC data portal (https://dcc.icgc.org/releases/PCAWG). Data for the IMmotion151 (NCT02420821, Motzer at al.)^30^, Wang et al.^40^, Durinck et al.^32^, Malouf et al.^99^, and Sato et al.^31^ cohorts were obtained from the corresponding studies. For the OncoPanel cohort, DNA extraction, sequencing, and mutation and copy number calling were performed as previously described for the OncoPanel gene panel assay^100^. The OncoPanel assay is an institutional analytic platform that is certified for clinical use and patient reporting under the Clinical Laboratory Improvement Amendments (CLIA) Act. The panel includes 275 to 447 cancer genes (versions 1 to 3 of the panel). Sample-level data for the OncoPanel cohort (mutations, gene-level CNA, and clinical metadata) are provided in **Supplementary Table 5**. The data types available for each dataset are illustrated in **Fig. 1C**, but not all data types were available for all samples in each cohort. The full list of samples used (including the data types available) and sequencing platform used for DNA-sequencing (WGS, WES, or panel) are provided in **Supplementary Table 1**.

### Analysis of mutation and copy number variants in genomic tRCC cohorts

Mutation calls (all aligned to human genome reference build hg19) were obtained as detailed above. Specifically, for the MSK cohort^15, 97^, WES-based calls were used where available and panel-based data were otherwise used for tRCC samples. For the TCGA cohort, the mc3 MAF calls^101^ (https://gdc.cancer.gov/about-data/publications/pancanatlas) were used. For the Durinck et al.^32^ and Malouf et al. cohort^99^, only samples from patients that had mutation calling based on matched normal sequencing were included. For the Sato et al. cohort^31^, only the WES calls were used. All mutations were annotated uniformly using Oncotator^102^ (except for the IMmotion151 cohort, for which a MAF was not available). In order to filter out potential germline mutations in the OncoPanel cohort, mutations present at an allelic frequency of 0.5% in one of the superpopulations from the 1000 Genomes Project^103^ (https://www.internationalgenome.org/data) were excluded from all downstream analyses. For the enrichment analyses, mutations were included if they were truncating (nonsense or splice site), insertions-deletions (indels), missense mutations, or *TERT* promoter mutations. For the IMmotion151 cohort, mutations were included if they were short-variants or truncating. The mutation load was computed as the number of all non-synonymous mutations per sample. The indel load was computed as the number of all indels per sample (either all indels or only frameshift indels). For the OncoPanel and MSK-IMPACT samples, the mutation and indel loads were normalized to the bait sets of the version of the panel used. The bait sets^104^ for OncoPanel were: v1, 0.753334 Megabases [Mb]; v2, 0.826167 Mb; and v3, 1.315078 Mb. For MSK-IMPACT, the bait sets were: IMPACT341, 0.896665; IMPACT410, 1.016478; and IMPACT468, 1.139322 Mb.

Gene-level copy number data calls were available for the MSK cohort^97^, IMmotion151 cohort^30^, OncoPanel cohort (**Supplementary Table 5b**), PCAWG (https://dcc.icgc.org/releases/PCAWG/consensus_cnv/GISTIC_analysis/ all_thresholded.by_genes.rmcnv.pt_170207.txt), and TCGA (http://firebrowse.org/; KIPAN dataset). For all gene-level analyses only focal events (deep deletions and high amplifications) were considered. As measures of the copy number alteration burden, the aneuploidy score and fraction genome altered were obtained for the TCGA^105^ and MSK^97^ cohorts, respectively. Arm-level calls were obtained for the TCGA cohort^105^.

### Genomic enrichment analyses

In order to account for the inherent differences between the included cohorts and to maximize the power of the study to detect differences in mutations and copy number alterations in tRCC versus other RCC histologies, a meta-analytic approach was adopted for all gene-level enrichment analyses, as has been done in prior studies^106, 107^. First, Fisher’s exact tests were used to evaluate the enrichment of gene alterations (mutations and copy number alterations separately) within each cohort (combined WES cohort, IMmotion151, PCAWG, OncoPanel, and MSK-IMPACT). For panel-based cohorts, this enrichment took into account the bait set of each version of the panel used for sequencing (i.e. a gene was counted as missing, and not non-mutated, if not included in the bait set of a version of the panel). The conditional maximal likelihood estimate of the odds ratio and its 95% confidence interval were computed using the fisher.test() function from the stats package in R. For each gene, we then obtained pooled estimates of the odds ratio and its 95% confidence interval using a random-effects model with the Paule-Mandel estimator for tau, with treatment arm continuity correction and Knapp-Hartung adjustment. The meta-analysis was performed using the metabin() function from the meta package in R^108–110^. The enrichment analysis was performed pairwise between tRCC and each comparator RCC histology separately (ccRCC, pRCC, and chRCC). Genes were included in the enrichment analysis if: (1) they were altered in at least two different cohorts; (2) alteration frequency in tRCC was 3% or more; and (3) were Tier 1 cancer genes as defined in the Cancer Gene Census (accessed on February 17 2021)^111^. Genes that had been previously reported to be significantly mutated in ccRCC, pRCC, and chRCC^27^ were also included in the analysis. For all analyses, samples that were originally part of the TCGA and PCAWG cohorts were only included in one of the two cohorts as part of the enrichment analyses (cohort assignment reported in **Supplementary Table 1**). The CoMut plot was generated using the CoMut package in Python^112^ and genes that were not assessed in specific samples (i.e. not included in the bait sets of the gene panel used) are shown as gray boxes; the corresponding alteration frequency (bar graph at the right-hand side of the CoMut) was adjusted accordingly and reflects only samples in which a particular gene was assessable for alteration. Arm-level comparisons (TCGA cohort) were performed pairwise with RCC histologies using Fisher’s exact tests. The mutation and indel loads, as well as the aneuploidy score and fraction genome altered, were compared pairwise with each RCC histology (ccRCC, pRCC, and chRCC) using Wilcoxon rank-sum tests.

### *MiT/TFE* fusion identification and characterization

Fusion calls were obtained as detailed under “Genomic tRCC cohorts” above. In particular, for the MSK cohort, determination of fusion partners was based on MSK-IMPACT and/or RNA-seq^15, 97^ and fusion breakpoints were based on MSK-IMPACT and available for a subset of samples^97^. For the OncoPanel cohort, fusion partners and breakpoints were based on an in-house fusion calling pipeline and were available for a subset of samples. For the TCGA, PCAWG, Wang et al., Sato et al., Durinck et al., and Malouf et al. cohorts, fusion partners were based on RNA-seq. Of those, the fusion breakpoints were available for the TCGA, PCAWG, Sato et al., and Durinck et al. cohorts. For the Malouf et al. cohort, fusion breakpoint locations were inferred based on the reported fusion breakpoint sequences using BLAT (https://genome.ucsc.edu/cgi-bin/hgBlat). All breakpoint locations were aligned to human genome reference build hg19, except for the TCGA breakpoints which had been originally mapped to hg38 and were converted to hg19, for the purposes of this analysis, using liftOver (https://genome.ucsc.edu/cgi-bin/hgLiftOver). The Circos Perl package^113^ was used to represent the chromosomal locations of fusions in a circos plot. The enrichr^114^ tool was used to evaluate enrichment of Gene Ontology (GO) terms among the *MiT/TFE* partner genes. In order to annotate the fusion protein products based on the breakpoints, breakpoints were first aligned to human genome GRCH37.p13 on NCBI Genome Data viewer. Functional domains were then annotated using UniPort Protein knowledgebase UniProtKB/Swiss-Prot And NCBI Conserved Domain Database^115^ (CDD v3.19). The presence of Prion-Like domains (PLD) was analyzed using Prion-Like Amino Acid Composition (PLAAC) web-based program^116^. Illustrations were made using Illustrator for Biological Sequences (IBS)^117^ version 1.0. Annotated functional domains with abbreviations are provided in **Supplementary table 2**.

### Cell lines

293T cells were obtained from the American Type Culture Collection. UOK109 and UOK146 cells were a kind gift of Dr. Marston Linehan (National Cancer Institute). FU-UR-1 cells were a kind gift of Dr. Masako Ishiguro (Fukuoka University School of Medicine). Cell lines were grown in base media of DMEM (293T, UOK109, UOK146) or DMEM/F12 (FU-UR-1), supplemented with 10% FBS, 100 U mL^-1^ penicillin, 100 μg mL^-1^ streptomycin, 2 mM L-glutamine, and 100 μg mL^-1^ Normocin (Invivogen).

### *TFE3* fusion-specific signature

For *TFE3* fusion overexpression experiments, 293T cells were seeded in 6-well plates at 2 x 10^5^ cells per well and after 24 hours were transfected with 500 ng of plasmids encoding *ASPSCR1-TFE3*, *NONO-TFE3*, *PRCC-TFE3*, *SFPQ-TFE3*, wild type (WT) *TFE3*, or an empty vector control (all in pLX313). All transfections were performed in three biological replicates. Cells were harvested 48 hours after transfection and total RNA was collected using the RNeasy Plus Mini Kit (QIAGEN, #74136). Sample concentrations were measured using a NanoDrop 8000 Spectrophotometer (Thermo Fisher Scientific) and sequencing libraries were prepared with poly(A) selection. Libraries were pooled and paired-end 150 bp RNA-sequencing was performed on an Illumina HiSeq. Paired-end sequencing reads were aligned to the human genome reference build hg38 using STAR v2.7.2^118^ and quantified using RSEM v1.3.2^119^. Transcripts were filtered based on read support (sum of expected read counts across three biological replicates > 30) prior to gene-level differential expression analysis using the voom transformation in limma v3.40.6^120^. Transcripts-per-million (TPMs) were used for visualization and clustering. Expected count and TPM matrices are provided in **Supplementary Table 3.**

In order to derive a transcriptional signature that is specific to the *TFE3* fusion, we performed differential gene expression of each of the fusion conditions (*ASPSCR1-TFE3*, *NONO-TFE3*, *PRCC-TFE3*, *SFPQ-TFE3*) versus the WT *TFE3* condition. Genes that were significantly upregulated (q<0.05 and log2(fold-change)>0) or significantly downregulated (q<0.05 and log2(fold-change)>0) across all four comparisons defined a *TFE3* fusion-specific signature (**Supplementary Table 4)**. In order to evaluate the relevance of the *in vitro*-derived signature to tRCC tumor samples, we performed clustering on 4 independent RNA-seq datasets that included tRCC samples. The normalized expression matrices used for clustering were those obtained from TCGA (https://gdc.cancer.gov/about-data/publications/pancanatlas), PCAWG (https://dcc.icgc.org/releases/PCAWG), IMmotion151, and Wang et al. as described under “Genomic tRCC cohorts” above. Clustering was performed in each dataset independently using the Heatmap function from the ComplexHeatmap^121^ package in R, using hierarchical clustering with ward.D2 as the clustering method and the Kendall correlation distance metric. The average intra-tRCC distance was used as a metric for density of clustering of tRCCs and was compared to the distance obtained from clustering using the 1000 most variable genes within each dataset (**Fig. S4**).

### Differential gene expression analysis

Pairwise differential gene expression analysis was performed between tRCC and each other sample type, within each dataset independently (TCGA, PCAWG, IMmotion151, Wang et al., and 293T cell line experiment). Differential gene analysis for the cell line experiment was performed as described above using the limma package. For the tumor datasets, differential gene expression was performed using pairwise Wilcoxon rank-sum tests. For all tests, the Benjamini-Hochberg correction was used to compute q-values and a q-value<0.05 was taken as statistically significant. In order to define a transcriptional signature specific to tRCC, an Upset plot was computed using the UpsetR package^122^. The 76 genes that were found to be significantly upregulated in 9 or more of the 14 pairwise comparisons were plotted in a heatmap (**Fig. 4c**), which included tRCC samples and comparator samples. Gene pathway annotations were obtained from enrichr^114^. Overlap between the NRF2 and MITF target genes was evaluated using a one-tailed hypergeometric test. In order to adjust for potential RNA-seq batch effects between datasets in visualization, gene expression was Z-scored within each dataset independently. For volcano plots, log2(fold-change) of the mean expression of genes in each group was used.

### Gene set enrichment analysis

Pre-ranked gene set enrichment analysis (GSEA) was performed pairwise between tRCC and each comparator, within each dataset independently, using the using -log10(q-value) signed by the sign of the log2(fold-change) of mean gene expression. GSEA was used on the Hallmark gene sets v7.1 from the Molecular Signatures Database (MSigDB)^123^ and a previously defined 55-gene NRF2 signature^124^. For the Hallmark analysis, the gene sets were ranked by the number of pairwise comparisons that had a normalized enrichment score (NES)>1 in tRCC vs the other comparators (with the top gene sets visualized as a dot plot) (**Fig. 4d**).

In addition, single sample GSEA (ssGSEA) scores were computed for the 55-gene NRF2 signature using the GSVA package^56^ in R to infer the level of activity of NRF2 in each sample. In order to adjust for potential RNA-seq batch effects in visualization, NRF2 signature scores were Z-scored within dataset prior to visualization as a waterfall plot (**Fig. 5a**). Comparison of ssGSEA scores between tumor types in the TCGA cohort was performed using Wilcoxon rank-sum tests. To examine the relationship of the NRF2 signature with survival outcomes, the NRF2 score was dichotomized at the median in each treatment arm of each cohort.

### Analysis of CTRP and DepMap datasets

RNAi genetic dependence scores were obtained from the DEMETER2 Data v6 dataset^125^, CRISPR genetic dependence scores were obtained from the CRISPR (Avana) Public 21Q1 dataset^126, 127^ and drug area under the curve (AUC) values were obtained from the CTRP v2.0 2015 CTD^2^ dataset^61, 128^. Cell lines were excluded if they had multiple AUC values for each drug. All datasets were downloaded from the DepMap Data Download Portal (https://depmap.org/portal/download/). NRF2 ssGSEA scores were calculated from the Broad Institute CCLE RNA-seq dataset. Expression values were upper quartile normalized prior to analysis. For each drug (or gene), drug AUCs (or gene dependence scores) were Z-scored and the NRF2 ssGSEA scores were Z-scored, amongst samples having data for both data types. Pearson’s correlation coefficient was used to assess the association between drug AUC Z-score and NRF2 ssGSEA Z-score as well as between gene dependence Z-score and NRF2 ssGSEA Z-score. For each correlation, t-statistics were computed (t = r•((n-2)/(1-r^2^))^0.5^), a two-tailed Student’s t-distribution was used to determine p-values, and q-values were computed using a Benjamini-Hochberg correction.

### Colony forming assays

shRNAs were cloned into a doxycycline-inducible lentiviral vector as previously described^129^. The indicated cell lines were transduced with lentivirus expressing doxycycline-inducible shRNA (shRNA target sequence: CCGGCATTTCACTAAACACAA) and selected with 500 μg/mL of G418 prior to seeding at equal densities with or without the addition of 1 μg/mL doxycycline. Cell densities ranged from 500-1500 cells per well of 12-well plate depending on the cell line. Fresh complete culture media with/without doxycycline was replaced every two days prior to fixation and staining with crystal violet after 12-20 days. Colony areas were quantified using Image J v1.53.

### Multiplex immunofluorescence and image analysis

Cluster of differentiation (CD8), programmed death 1 (PD1), T cell immunoglobulin and mucin domain-3 (TIM3), and Lymphocyte-activation gene 3 (LAG3) multiplex immunofluorescence (IF) was performed as previously described^66^. Briefly, we used the Perkin Elmer Opal tyramide signal system on a Bond RX Autostainer (Leica Biosystems). The anti-CD8 antibody (1:5,000, C8/144B, mouse monoclonal antibody, Agilent) was detected using the Opal 520 fluorophore (1:150, FITC); the anti-TIM3 antibody (1:1,000, AF2365 goat monoclonal antibody, R&D Systems) was detected using the Opal 540 fluorophore (1:50, Cy3); the anti-LAG3 antibody (1/10,000, 17B4 mouse monoclonal antibody, LifeSpan Biosciences) was detected using the Opal 560 fluorophore (1:150, Texas Red); the validated anti-PD-1 antibody (1:5,000, EH33 mouse monoclonal antibody, Dr. Freeman laboratory, Dana-Farber Cancer Institute, Boston, MA) was detected using the Opal 690 fluorophore (1:50, Cy5). Whole slide images were acquired at 10x using the Vectra 3 automated quantitative pathology imaging system (PerkinElmer). Subsequently, at least 5 stamps of 931×698 um were selected per slide in areas of high immune infiltration (hotspots) using Perkin Elmer Phenochart v 1.0 software. Each stamp was then acquired at 20x using the Vectra 3. Inform 2.2 software was then used in order to deconvolute the multispectral images, as previously described^71^. Hotspot deconvoluted images in .tiff format were uploaded into Indica Lab HALO platform version 3.0. For each hotspot, the tumor area was manually annotated by a pathologist (TD). CD8 cells were phenotyped according to the expression of PD1, TIM3 and LAG3 using the Indica Lab High-Plex FL v2.0 module, using DAPI-based nuclear segmentation and detection of FITC (CD8), Cy3 (TIM3), Texas Red (LAG3), Cy5 (PD1) positive cells by adapting a dye cytoplasm positive threshold for each slide. A unique algorithm was created for each whole slide, and each group of hotspots and its accuracy was validated through visual inspection by two pathologists (TD, SS). Sample-level results of the multiplex immunofluorescence analysis are provided in **Supplementary Table 6**. Comparisons between tRCC (n= 11), ccRCC (n= 11), and normal (ccRCC adjacent, (n= 10)) were performed using Wilcoxon rank-sum tests. All tRCC samples were either (1) *TFE3* FISH positive or (2) positive TFE3 test by IHC along with a strongly suggestive clinico-pathologic history and no FISH testing results available (missing). For each T cell subset, T cell subset density was calculated as the number of T cells per mm^2^. Percentage of a T cell subset was defined as the density of the T cell subset divided by the density of CD8^+^ T cells in the sample.

### Immune deconvolution and immune analyses

CIBERSORTx^70^ (Job type: “Impute cell fractions”), in absolute mode, with B mode batch correction, with quantile normalization disabled, and in 1000 permutations was used on the LM22 signature in order to infer the immune cell composition of samples from RNA-seq in the TCGA cohort. All samples which had a p-value for deconvolution >0.05 were considered to have failed deconvolution and were therefore discarded from all downstream analyses. Relative cell proportions were obtained by normalizing the CIBERSORTx output to the sample-level sum of cell counts (in order to obtain percentages of immune infiltration). Purity estimates for the TCGA cohort were obtained for the TCGA cohort from the Taylor et al. study^105^. CD8^+^ T cell density and purity were compared pairwise between tRCC and each other RCC histology (ccRCC, pRCC, and chRCC) using Wilcoxon rank-sum tests. Sample-level PD-L1 protein expression by IHC on tumor-infiltrating immune cells (PD-L1≥ 1%) for the IMmotion151 trial were obtained from the Motzer et al. study^30^ and compared using a Fisher’s exact test between tRCC and ccRCC.

### Clinical and survival analyses

Tumor stage was obtained from Genomic Data Commons (https://gdc.cancer.gov/about-data/publications/pancanatlas) for the TCGA cohort and was defined using American Joint Committee on Cancer (AJCC) 8^th^ edition for the IMDC and Harvard cohorts. IMDC risk groups (a previously validated prognostic model for patients with metastatic RCC) were defined as previously described^130^. Tumor stage (I/II vs III/IV), IMDC risk groups (favorable, intermediate, poor), and sex were compared pairwise between tRCC and each other RCC histology (ccRCC, pRCC, and chRCC) using Fisher’s exact test. Age at initial RCC diagnosis was compared between tRCC and each other RCC histology (ccRCC, pRCC, and chRCC) using Wilcoxon rank-sum tests. Sankey diagrams for the Harvard and IMDC cohorts were computed using the ggalluvial package in R.

For all survival endpoints, the Kaplan-Meier method was used to summarize survival distributions. For the TCGA cohort, progression-free interval (PFI) was defined as the period from the date of diagnosis until the date of the first occurrence of a new tumor event (includes disease progression, locoregional recurrence, distant metastasis, new primary tumor, or death with tumor). Patients were censored if they were alive without any of these events at last follow-up or had died without tumor^131^. Overall survival (OS) was defined as the period from the start of systemic therapy until death. Patients were censored if they were alive at last follow-up. Time-to-treatment failure (TTF) was defined from the start of the line of systemic therapy to the end of that line of therapy or death from any cause. Since assessment of responses in retrospective cohorts (Harvard and IMDC cohorts) was not subject to radiological review specifically for the purpose of this study, responses were defined based on RECIST v1.1 criteria^132^ as available by retrospective review. Patients were censored if they were alive and still on the line of therapy at last follow-up. Progression-free survival (PFS) was defined (in the CheckMate and IMmotion151 cohorts) from the time of randomization or start of first dose until disease progression or death. Patients were censored if they were alive at last follow-up. For all survival endpoints, pairwise comparisons were performed using log-rank tests. In the IMmotion151 cohort, a Cox model that included an interaction term (histology-by-treatment arm) was used to evaluate the difference in the extent of benefit derived with atezolizumab + bevacizumab versus sunitinib in tRCC versus ccRCC. In the Harvard/IMDC pooled cohort, all patients who got ICI-based therapies were included in the ICI group. If patients received multiple lines of ICI-based therapies, the first ICI-based regimen was used for the analysis of clinical outcomes. All other patients had received TKIs and were assigned to the TKI group. If patients received multiple lines of TKIs, the first TKI regimen was used for the analysis of clinical outcomes.

Clinical benefit (CB) was defined as an objective response (complete response or partial response) or stable disease with PFS of at least 6 months. No clinical benefit (NCB) was defined as progressive disease with PFS less than 3 months. All other patients (not meeting criteria for CB or NCB) were classified as having intermediate clinical benefit (ICB). In the IMmotion151 cohort, rates of CB and NCB were compared between the atezolizumab + bevacizumab and sunitinib arms, in patients with tRCC and ccRCC separately, using Fisher’s exact test.

### Statistics

All downstream analyses were done using R v3.6.1, Python v3.8.5 (on Spyder v4.1.5), Circos v0.69.9, or GraphPad PRISM 9. For boxplots, the upper and lower hinges represent the 75^th^ and 25^th^ percentiles, respectively. The whiskers extend in both directions until the largest or lowest value not further than 1.5 times the interquartile range from the corresponding hinge. All tests were two-tailed (unless otherwise specified) and considered statistically significant if p< 0.05 or q< 0.05.

### Data availability statement

All relevant data are available from the authors and/or are included with the manuscript. The list of samples used (including the data types available) and sequencing platform used for DNA-sequencing (WGS, WES, or panel) are provided in **Supplementary Table 1**. The expression matrix (RSEM expected counts and TPMs) derived from the RNA-sequencing of the cell lines in the *in vitro* experiment represented in **Figure 4a** is provided in **Supplementary Table 3.** For the OncoPanel cohort, sample-level data (mutation, copy number, and clinical metadata) are provided in **Supplementary Table 5.** Sample-level data from the multiparametric immunofluorescence cohort are provided in **Supplementary Table 6.**

### Code availability statement

Algorithms used for data analysis are all publicly available from the indicated references in the **Methods** section. Any other queries about the custom code used in this study should be directed to the corresponding authors of this study.

## Acknowledgements

We thank the OncoPanel study team and the patients who contributed their data to research and participated in clinical trials. We thank all contributors to The International Metastatic Renal-Cell Carcinoma Database Consortium for their data contributions. This work was supported in part by The Friends of Dana-Farber (S.R.V.), Claudia Adams Barr Program for Innovative Cancer Research (S.R.V.), Clinician-Scientist Development Award from the Doris Duke Charitable Foundation (S.R.V.), Department of Defense Kidney Cancer Research Program (W81XWH-19-1-0815) (S.R.V.), an Independent Investigator Grant from the Rally Foundation for Childhood Cancer Research (S.R.V), and the Dana-Farber / Harvard Cancer Center Kidney Cancer SPORE (P50-CA101942) (D.A.B., D.F.M., S.S., T.K.C., S.R.V.). T.K.C. is supported by the Kohlberg Chair at Harvard Medical School and the Trust Family, Michael Brigham, and Loker Pinard Funds for Kidney Cancer Research at DFCI. J.N. acknowledges support by NIH F31 CA250136. N.I.V. is supported by Damon Runyon Physician-Scientist Training Award, Conquer Cancer Foundation YIA, SITC Genentech Award. D.A.B. is supported by the Dana-Farber / Harvard Cancer Center Kidney Cancer SPORE Career Enhancement Program (NCI P50CA101942-15), DoD CDMRP (KC170216, KC190130), and the DoD Academy of Kidney Cancer Investigators (KC190128).

## Author contributions

Conception and Design: Ziad Bakouny, Eliezer M. Van Allen, Toni K. Choueiri, Srinivas R. Viswanathan Provision of study material or patients: Ziad Bakouny, Praful Ravi, Xin Gao, David A. Braun, Eliezer M. Van Allen, Toni K. Choueiri, Srinivas R. Viswanathan

Collection and assembly of data: Ziad Bakouny, Praful Ravi, Jiao Li, Stephen Tang, Thomas Denize, Emma R. Garner, Laure Hirsch, John A. Steinharter, Gabrielle Bouchard, Emily Walton, Destiny West, Chris Labaki, Shaan Dudani, Chun-Loo Gan, Vidyalakshmi Sethunath, Filipe LF. Carvalho, Michelle S. Hirsch, Gwo-Shu Mary Lee, Bradley A. McGregor, Steven L. Chang, Adam S. Feldman, Catherine J. Wu, David F. McDermott, Daniel Y.C. Heng, Eliezer M. Van Allen, Toni K. Choueiri, Srinivas R. Viswanathan

Data Analysis and Interpretation: Ziad Bakouny, Ananthan Sadagopan, Nebiyou Y. Metaferia, Shatha AbuHammad, Stephen Tang, Thomas Denize, David A. Braun, Alma Imamovic, Cora Ricker, Natalie I. Vokes, Jackson Nyman, Jihye Park, Rizwan Haq, Eliezer M. Van Allen, Toni K. Choueiri, Srinivas R. Viswanathan

Manuscript writing and revision: All authors

Final approval of manuscript: All authors

Accountable for all aspects of work: All authors

---

**Fig. S1.**
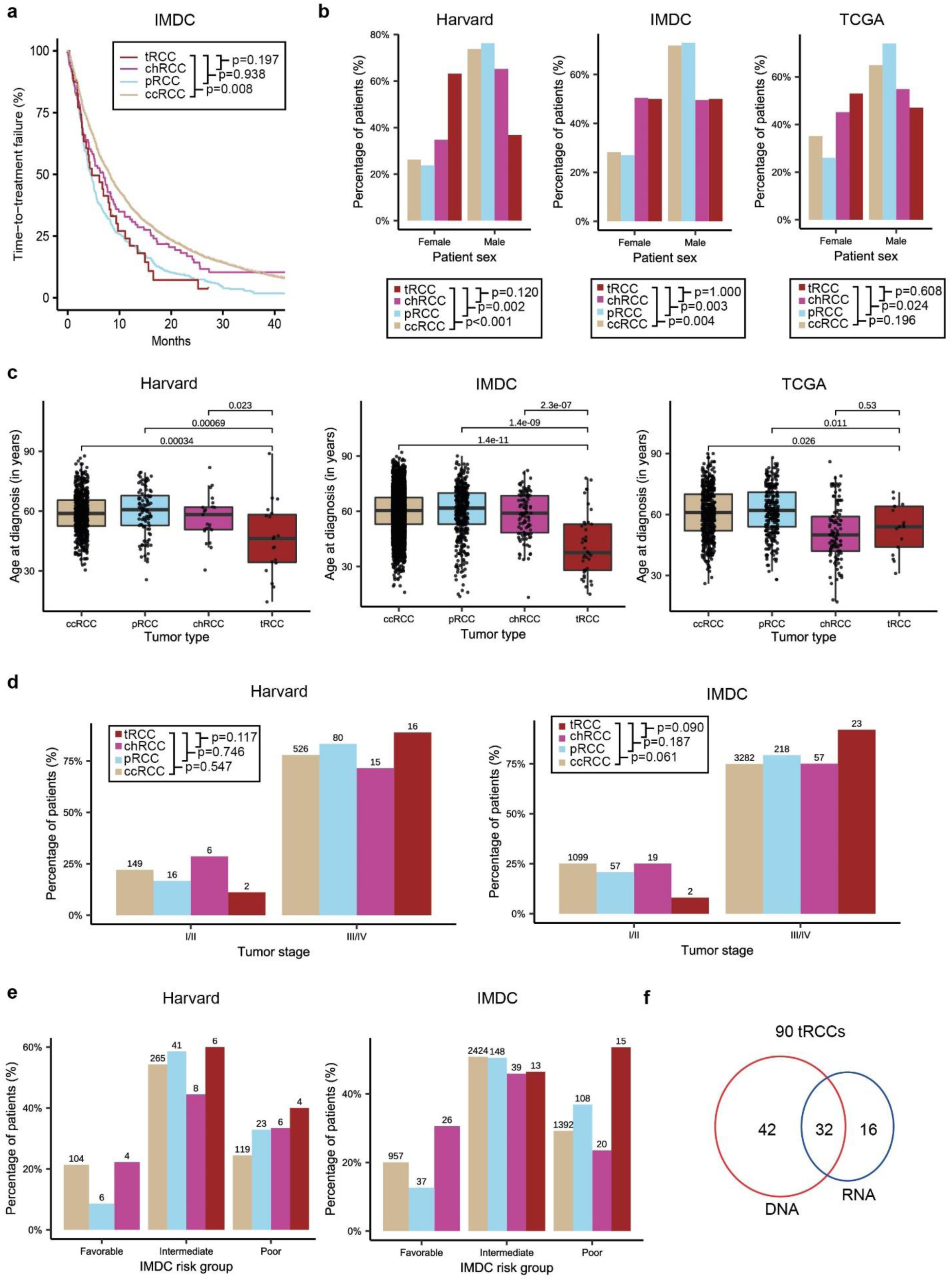
Clinical features of tRCC. **a**, Kaplan-Meier curves for time-to treatment-failure in metastatic ccRCC, pRCC, chrRCC, and tRCC from patients in the IMDC cohort. **b**, Proportion of male and female ccRCC, pRCC, chrRCC, and tRCC cases in the Harvard, IMDC, and TCGA cohorts. **c**, Age distribution of tRCC, ccRCC, chRCC, and pRCC cases in the Harvard, IMDC, and TCGA datasets. **d** Distribution stage at diagnosis among ccRCC, pRCC, chrRCC, and tRCC patients in the Harvard and IMDC cohorts. **e**, Distribution of IMDC risk group at start of first-line of systemic therapy among ccRCC, pRCC, chrRCC, and tRCC patients in the Harvard and IMDC cohorts. **f**, Number of tRCC samples with DNA sequencing (WGS, WES, or Panel sequencing), RNA sequencing, or both data types, available for analysis across all NGS data sets.

**Fig. S2.**
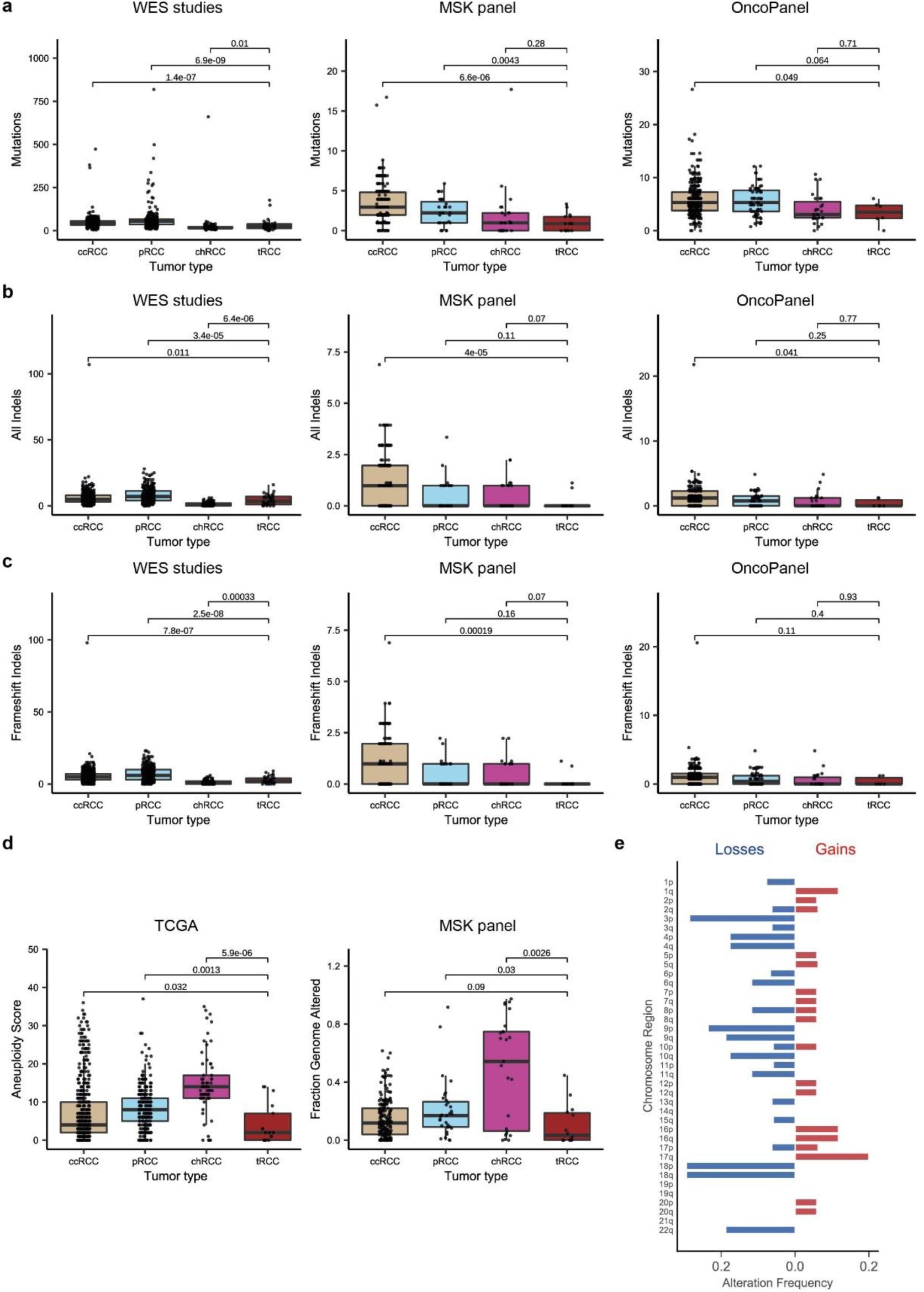
Genomic features of tRCC as compared with other RCC subtypes. **a**, Number of mutations per sample in tRCC versus other RCC histologies in the TCGA, MSK, and OncoPanel cohorts. **b**, Number of indels per sample in tRCC versus other RCC histologies in the TCGA, MSK, and OncoPanel cohorts. **c**, Number of frameshift indels per sample in tRCC versus other RCC histologies in the TCGA, MSK, and OncoPanel cohorts. In **a-c**, for the OncoPanel and MSK cohorts, the numbers of mutations and indels were normalized to the bait set of each version of each panel (**Methods**) **d**, *Left*, Aneuploidy score^34^ in tRCC versus other RCC histologies in the TCGA cohort. *Right*, Fraction of genome altered in tRCC versus other RCC histologies in the MSK cohort. **e**, Frequency of arm-level copy number alterations among tRCC samples in the TCGA cohort^34^.

**Fig. S3.**
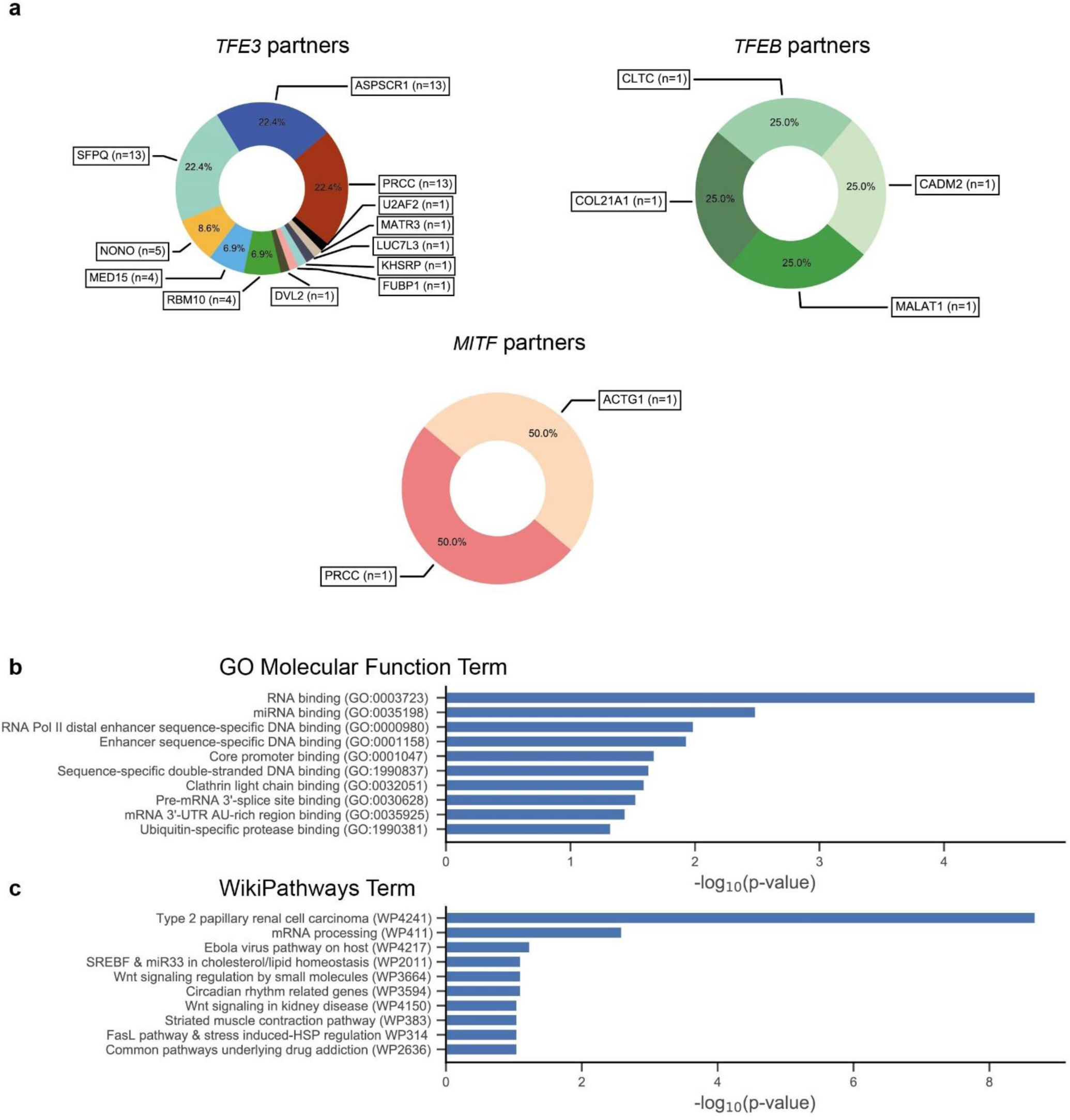
| Characterization of *MiT/TFE* fusion partners. **a,** Frequency of various partner genes observed to fuse with *TFE3*, *TFEB*, or *MITF* across all datasets. **b-c,** Terms enriched amongst *MiT/TFE* fusion partner genes using either the GO Molecular Function **(b)** or Wikipathways **(c)** databases.

**Fig. S4.**
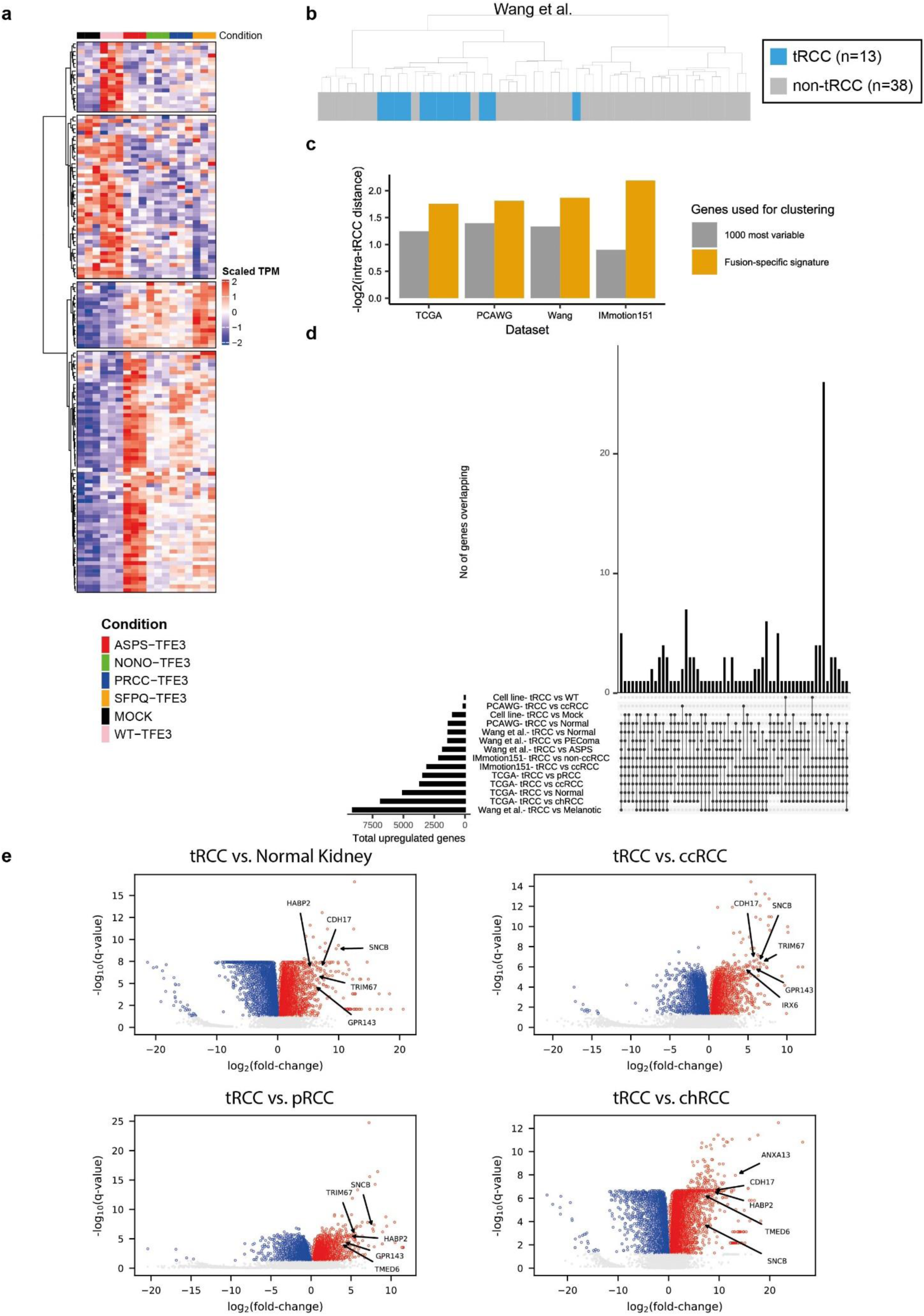
Transcriptional features of tRCC. **a,** Expression of genes included in the *in vitro*-derived *TFE3* fusion-specific gene signature. **b**, Hierarchical clustering of RNA-Seq data^134^ based on fusion-specific gene signature. **c**, Quality of clustering (based on -log2(intra-tRCC distance)) in the TCGA, PCAWG, Wang et al., or IMmotion151 datasets using either 1000 most variable genes (grey) or the fusion-specific gene signature (orange). **d**, Upset plot showing overlap of upregulated (q<0.05) genes in tRCC versus other sample types in each of the datasets analyzed. **e**, Volcano plots showing differentially expressed genes in tRCC samples versus normal kidney, ccRCC, pRCC, and chrRCC in the TCGA cohort. Labels indicate selected genes that emerged as commonly upregulated in tRCC versus other sample types (Figures 4c and S4c) across all datasets analyzed.

**Fig. S5.**
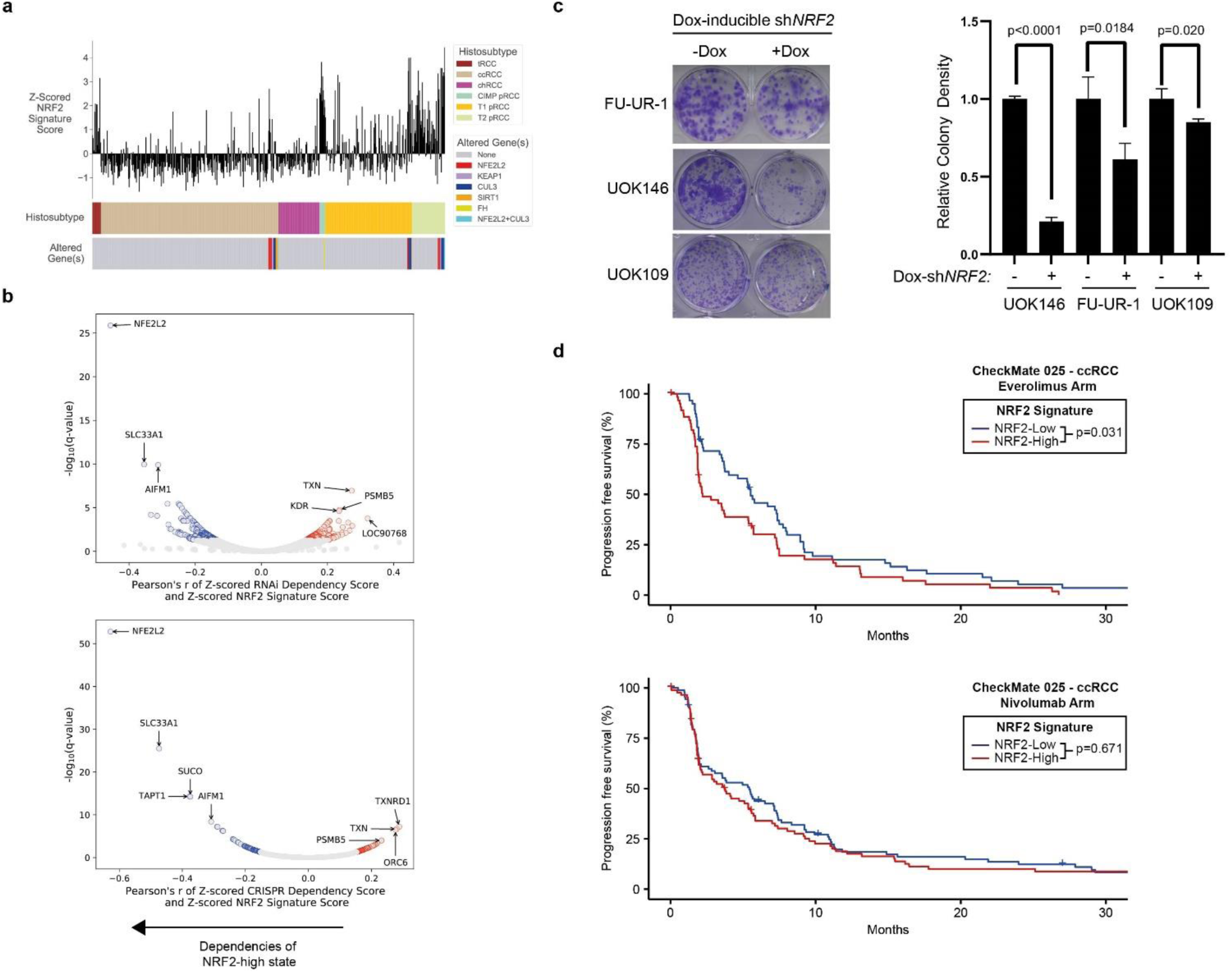
Activation of the NRF2 pathway in tRCC. **a**, NRF2 signature score in tRCC samples compared with ccRCC, pRCC, or chrRCC samples from the TCGA effort. Papillary RCC subtypes are annotated as previously described^26, 27^. Somatic alterations in the NRF2 pathway genes are indicated on the bottom track. **b**, Volcano plot displaying gene dependencies correlated to high NRF2 score in the DepMap RNAi (top) and CRISPR (bottom) datasets. **c**, Colony-forming assay in three tRCC cell lines (FU-UR1, UOK109, UOK146) transduced with a lentiviral doxycycline-inducible shRNA targeting *NRF2*. Quantification represents mean +/-s.d. for n=3 independent replicates. **d**, Progression-free survival curves for ccRCC patients with high (red) or low (blue) NRF2 signature score treated with either everolimus (top) or nivolumab (bottom) in the CheckMate cohort. NRF2 signature score was dichotomized at the median in each arm.

**Fig. S6.**
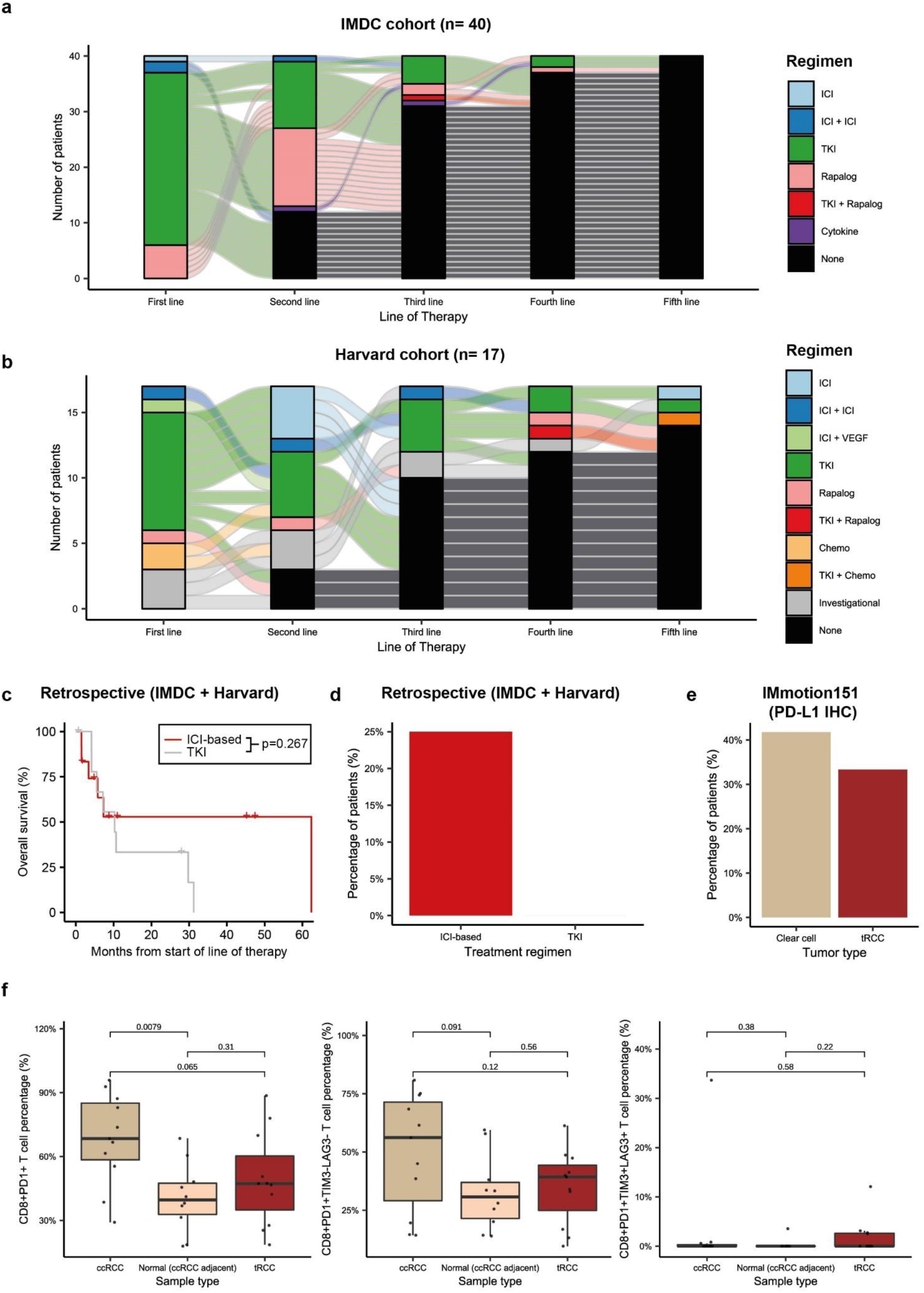
Immunogenomic features and treatment patterns in tRCC. **a,** Sankey flow diagram showing lines of systemic treatment received by patients with metastatic tRCC in the retrospective IMDC cohort (n=40). **b**, Sankey flow diagram showing lines of systemic treatment received by patients with metastatic tRCC in the retrospective Harvard cohort (n=17). **c**, Kaplan-Meier curves for overall survival in metastatic tRCC patients who received ICI-based (n=12) or tyrosine kinase inhibitor (TKI, n=10) regimens in the combined Harvard + IMDC retrospective cohort. **d**, Percentage of tRCC patients showing a response to either immune checkpoint inhibitor (ICI-based) or tyrosine kinase inhibitor (sunitinib and pazopanib) in the com bined IMDC and Harvard retrospective cohorts. **e**, PD-L1 protein expression on infiltrating immune cells (PD-L1≥ 1%) in tRCC (n=15) and ccRCC (n=797) in the IMmotion151 cohort. **f,** Quantification of percentage of CD8^+^PD1^+^ T-cells (left), percentage of CD8^+^PD1^+^TIM3^-^LAG3^-^ T cells (middle), and percentage of CD8^+^PD1^+^TIM3^+^LAG3^+^ T cells (right) in tRCC (n=11), ccRCC (n=11), and adjacent normal tissue (from ccRCC cases, n= 10) analyzed by multiparametric immunofluorescence.

**Supplementary Table 1**: List of samples in the NGS datasets included in the analysis.

**Supplementary Table 2:** List and legend of functional domains used in the annotation of *MiT/TFE* and partners genes in Figures 3d-e.

**Supplementary Table 3:** RSEM expected counts (**Supplementary Table 3a**) and transcript-per-million (TPM; **Supplementary Table 3b**) derived from the RNA-sequencing of the cell lines in the *in vitro* experiment represented in Figure 4a.

**Supplementary Table 4:** List of genes that are in the *TFE3*-fusion-specific transcriptional signature developed in Figure 4a.

**Supplementary Table 5:** Sample-level MAF (**Supplementary Table 5a**) and gene-level copy number (**Supplementary Table 5b**) data for the OncoPanel cohort.

**Supplementary Table 6:** Sample-level data for the multiparametric immunofluorescence cohort.

